# Drivers of mosquito free-flight and resting behavior indoors

**DOI:** 10.64898/2026.02.11.705377

**Authors:** David Jimenez-Vallejo, Gabriela Gonzalez-Olvera, Azael Che-Mendoza, Anuar Medina-Barreiro, Felipe Del Castillo-Centeno, Storm A. Crews, Guillaume Bastille-Russeau, Pablo Manrique-Saide, Gonzalo Vazquez-Prokopec

## Abstract

Mosquito flight and resting behaviors mediate pathogen transmission and vector control success, yet their environmental and genetic determinants remain poorly understood. We combined novel high-resolution 3D tracking with factorial semi-field experiments in Mérida, Mexico, to quantify how endogenous traits (strain, sex, physiological state) and exogenous conditions (microclimate, visual cues) influence the flight and resting behavior of Aedes aegypti, the primary vector of Dengue, Zika, and Chikungunya. Mosquitoes with a Wild-type genetic background flew and rested at lower heights than mosquitoes from with a laboratory strain background. Such difference was not explained by microclimate, with progeny from reciprocal cross experiments demonstrating wild-type-like behavior, which suggest a potentially heritable behavioral divergence. Microclimatic (temperature and relative humidity) gradients differed across time and height, with estimated Vapor Pressure Deficits (VPDs) indicating that mosquitoes adjusted flight height to minimize desiccation risk. However, this microclimatic influence was overridden by presence of black surfaces, which strongly attracted mosquitoes to rest, even if such resting heights had unfavorable conditions. These findings reveal a trade-off between visual, genetic, and microclimatic drivers of behavior which have important influence in the design and impact of vector control interventions.

**Significance Statement:** This study integrates high-resolution, three-dimensional mosquito tracking with analogue sticky traps and mathematical modeling to understand innate free-flight and resting behavior of indoor-dwelling mosquitoes. Factorial semi-field experiments conducted in Merida, Mexico, demonstrate that wild and laboratory mosquito strains differ markedly in flight and resting preferences. By quantifying microclimatic gradients and vapor-pressure deficits, we show mosquitoes may dynamically adjust their vertical position to minimize desiccation risk, however, presence of strong visual cues can override these microclimatic constraints. Together, we demonstrate a mechanistic framework that links genetics, physiology, and environment to explain Ae. aegypti behavior indoors.

## Introduction

Insect movement, particularly flight, is governed by a complex repertoire of sensory inputs, environmental stimuli, and internal physiological processes that are often difficult to decode ^1–9^. Particularly for mosquitoes, the relative importance of such mechanisms is fundamental to understanding behaviors such as host-seeking, mating, oviposition, and resting, which directly influence survival, reproduction, and pathogen transmission potential ^4,10–13^. Recent advancements in high-resolution tracking technologies have provided new opportunities to investigate mosquito behavioral dynamics under controlled laboratory conditions ^14–17^. Remarkably, one area that has lagged in research has been the study of mosquito free flying behavior outside of controlled laboratory conditions. Addressing this knowledge gap will enable better understanding of mosquito biology and the development of novel vector control strategies ^1,14,15,18–24^.

*Aedes aegypti* transmits Dengue, Zika, and Chikungunya primarily in tropical and sub-tropical regions ^25–29^. Although the species has its origin from a Sub-Saharan forest tree-hole ancestor, evolution drove *Ae. aegypti* to become a specialist by developing a strong preference to bite humans and breed within human urban environments ^30^. Given the availability of human hosts inside dwelling, females can easily obtain a blood-meal and complete their gonotrophic cycle indoors without needing to disperse long distances, often within 100-250 meters ^31–33^. It is important to note that dispersal (whether to find a blood-meal, a mate, or a resting site) only represents a narrow percentage of a mosquito’s entire time-activity budget ^34^. Direct observations and trap data have shown that *Ae. aegypti* partitions its diel cycle between two active periods (at dawn and at dusk) and resting periods ^4,6,23,35–37^ These resting periods can function to conserve energy or set aside time to other critical physiological (blood-digestion, oogenesis) and neurological (sleep) processes in times when other behaviors may be less efficient or of higher risk ^38–43^. Thus, in addition to flight, resting behavior is likely to be important for the survivorship, reproductive success, and disease dynamics of *Ae aegypti* ^34,40,41,44–46^.

Early observational studies of free-flying *Ae. aegypti* indoors provide evidence for a higher abundance of mosquitoes inside bedrooms, and a very particular preference to rest at heights below 1.5 meters ^47–51^ . Videography conducted in an experimental room in Brazil showed *Ae. aegypti* adults visit a narrow area at the lowest level of walls (0 – 20cm), suggesting that lower ambient temperatures and color (specifically, black) modulate such height preference. In addition to these height preferences, other research has provided strong evidence for *Ae. aegypti* to rest on dark objects/surfaces and under furniture (e.g. shadows), likely for camouflage ^47,48,50–53^.

Important technological advancements over the last few years have facilitated the development of automated tracking systems capable of recording multiple individuals over prolonged periods of time and of obtaining corresponding 3D reconstructions of flight paths ^1,7,8,15,16,18,34,51,54–56^. As stereoscopic tracking tools become more widely available, improved methods to analyze complex 3D data on mosquito flight paths are needed ^34^. Simple metrics such as mean flight height or track length may miss complex behaviors or even obscure relationships due to multiple behavioral phenotypes occurring within the same population. Such relationships may only be apparent after tracking hundreds or thousands of individuals, which is currently methodologically difficult. More robust quantitative methods (emerging from movement ecology literature) have been recently developed to address the challenge of animal movement in 3D, extending random walks, state-space models and machine learning algorithms to the third dimension^57^. Specifically for mosquitoes, various 2D and 3D models have been introduced to study flight^54,58^ without empirical linkage of models to observations on free-flying individuals. Better understanding the drivers of mosquito flight and fine-scale spatial arrangement and resting may provide important insights on behaviors that may lead to the success or failure of interventions, or to increased pathogen transmission risk ^34^.

Given the complexity of behavioral and physiological repertoires involved, both endogenous (e.g., sex, physiological state, genetic variability) and exogenous (e.g., resting sites, presence of a host, microclimate) factors may be important in determining the flight dynamics and resting preferences of *Ae. aegypti* in the field. Moreover, the strong preference in resting height and locations found across global *Ae. aegypti* populations cannot rule out interaction effects of environmental stimuli with the mosquito’s genetic background (GxE) on resting and flight preferences. Evidence of local adaptation at fine spatial scales to climate and vegetation has been described for *Ae. aegypti*^59^ yet translating such findings to behavioral traits has not been pursued. While evidence of sex-specific genetic control of flight through muscle development exist^60^, other genetic aspects related to flight and resting remain unexplored.

This study integrates experimental and theoretical frameworks to study the complex repertoire of factors influencing *Ae. aegypti* flight patterns and resting site selection indoors. By conducting a factorial design in experimental huts located in Merida, Mexico, which included direct tracking of 3D mosquito flight and location-specific resting, we evaluated the impact of endogenous and exogenous factors on the innate flight and resting behavior of *Ae. aegypti*. We hypothesized that flight and resting are the outcome of complex interactions between mosquito genetic background, microclimate, and the presence and type of resting surfaces, which combined guide a mosquito’s decision of where to fly and rest. A better understanding of the mechanisms and factors that drive *Ae. aegypti* to fly to certain areas or rest in specific locations could lead to better contextualize vector control interventions such as rear-and-release, residual insecticide applications or the deployment of spatial emanators indoors ^34^.

## Results and Discussion

Results are focused on unfed females, given their role in pathogen transmission, with comparative findings for males and blood-fed females presented in a separate section (see ‘Patterns between sex and physiological states’).

### Innate *Aedes aegypti* free-flight and resting

To quantify the innate pattern of wild-type (WT) *Ae. aegypti* flight, we used a 3D stereoscopic mapping tool within an experimental hut located in Merida, Mexico (‘Experimental Setup’ in Methods). From three 2-h experiments tracking 100 free-flying adult *Ae. aegypti* each, and with no resting stimuli, a total of 74,850 x,y,z coordinates from a total of 50 tracks lasting more than 1 minute (Fig. 1a) were analyzed. Turning angles were small (70% of the distribution <25°), indicating the majority of movements had smooth directional changes (Fig. 1b). Flight speed showed a significant linear increase with total distance traveled by each mosquito (Fig. 1c), indicating that mosquitoes traveling longer distances tended to fly faster, perhaps straighter. The fractal dimension (D, see Methods) evidenced such specific mix of behaviors (Fig. 1d): mosquitoes with D∼1 tended to fly linearly and make few turns whereas mosquitoes with D > 1.2 had more exploratory (tortuous) flight patterns (Fig. S4 shows sample tracks for each behavior). In the absence of stimuli, most mosquito tracks and their corresponding height coordinates tended to stay within the same height, changing slightly upwards or downwards but not towards any predominant direction (Fig. 1e).

**Figure 1.**
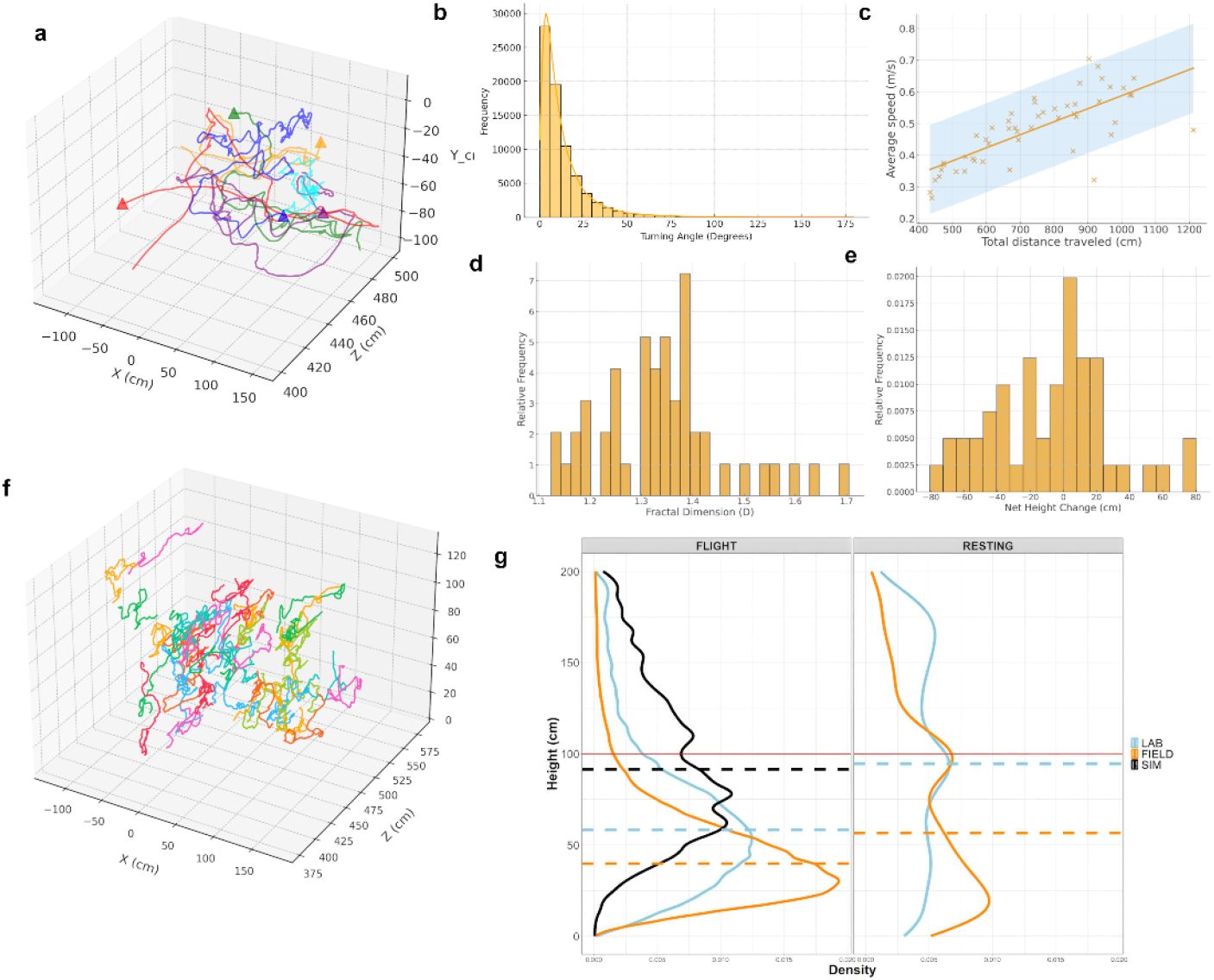
Innate free-flying and resting parameters of *Ae. aegypti* females. **a** sample of six mosquito tracks measured with the PFMD (triangles indicate the end-point of each track). **b** Distribution of turning angles for a total of 50 tracks which comprised ∼74000 observations (0 in vertical axis indicates the room’s 1m vertical point). **c** Relationship between the total distance traveled (length of track) and the average speed, showing a power-law fit with 95%CI. **d** Distribution of D (Fractal dimension) across the 50 tracks. When D∼1 movement is nearly straight, whereas for D∼1.5 more complex (tortuous) movements are implied. **e** Distribution of net height change (cm) across the 50 tracks. **f** Distributions of turning angles and step lengths were used to parameterize a random walk model simulating mosquito dispersal under no stimuli (see Methods, ‘Modeling mosquito random flight’). A random sample of 50 simulated tracks is shown. **g** Left: Cumulative density curves for flight height (*y*-coordinates) across all mosquito tracks recorded for lab (skyblue) and field (orange) mosquitoes, compared to density of flight height from simulated random tracks (black). Right: Cumulative density curves for resting heights from lab (skyblue) and field (orange) mosquitoes captured by sticky traps. In both, the dashed lines represent the median of each density curve.

Data from innate (non-stimulated) behavior was used to parameterize a 3D model of *Ae. aegypti* random flight (Fig. 1f, code available in Supplemental Materials). Unlike previous empirical descriptions of mosquito flight ^2,7,15,61^, the use of the random model provided a more reliable benchmark to compare behaviors. Assuming the mosquitoes move at random allowed hypothesis testing by fitting generalized linear mixed models (GLMMs) comparing observed and expected behaviors. Compared to the expectation of random flight, WT and Lab strains showed markedly reduced flight height (average track height 169.38, 36.01, and 52.27 cm, respectively; Fig 1g). Strain differences in track height were significant between random and WT(*β* = -3.15 ± 0.15, *z* = -20.37, *p* < 0.0001), and between random and Lab (*β* = -2.88 ± 0.15, *z* = -18.66, *p* < 0.0001, Table S1). Comparing between mosquito genetic backgrounds (i.e. excluding simulated null trajectories), WT flew significantly lower than the Lab strain (36.01cm vs 52.27cm, respectively; *β* = -0.342 ± 0.15, *z* = -2.224, *p* = 0.0261).

Resting heights, measured with four 30×200cm, non-attracting (i.e. white) sticky traps (see ‘Experimental Setup’ in Methods) showed a similar pattern as flight track height (Fig. 1f). Overall, WT mosquitoes rested significantly lower than Lab strain, at an average height of 34.2 cm versus 50.0cm, respectively. Fitting a GLMM to this data, a significant effect of strain on resting height (*β* = -1.037 ± 0.46, *z* = -4.46, *p* < 0.0001, Table S1) was observed. Comparing these findings to the flight averages, mosquitoes from both strains were, on average, choosing to rest at the same heights that they preferred to fly at (Fig 1g). However, WT mosquitoes showed a stronger preference for flying and resting at low heights than did the Lab strain. Lab strains like CDC-Rockefeller, which was used in this study, have high levels of inbreeding and low genetic diversity^62^. Finding such a marked statistical difference in flight and, more strongly, resting height in the absence of stimuli may be indicative of potential diverging behavioral preferences between strains. In the laboratory, mosquitoes are exposed to constant temperatures and flight environments. Given Rockefeller has been in the lab for at least a century^63^, the possibility for a genetic signal explaining differing flight and resting behaviors between strains cannot be ruled out.

### Exogenous factors influencing mosquito free-flight and resting: interplay of temperature, relative humidity, and mosquito flight and resting

Temperature and water availability are two of the most important exogenous (abiotic) variables influencing mosquito populations and pathogen amplification ^44,45,64–70^. We determined if a microclimatic preference may be driving the flight and resting preferences of WT *Ae. aegypti* females in the absence of attractive surfaces by capitalizing on the observed temperature and relative humidity gradients found within the experimental hut system (Figure S5, SI text). Generalized Additive Models (GAMs) (see ‘Statistical Analyses’ in Methods) revealed a significant non-linear interaction between temperature and relative humidity, influencing both flight height (flight model: edf = 7.99, X^2^ = 7590, *p* < 0.00001, Figure 2A) and resting height (resting model: edf = 3.864, X^2^ = 21.25, *p* = 0.00037; Figure 2B). Due to the difficulty of separating the behavioral effect of temperature and relative humidity, we characterized indoor microclimates by estimating vapor pressure deficit (VPD), which is considered a more biologically relevant metric for mosquitoes ^65,71,72^. VPD considers both temperature and relative humidity, providing a measurement of the difference between the amount of moisture the air *can* hold and the amount it *currently* holds, and thus higher VPD values indicate drier air (or increased desiccation pressure). Across all replicates, VPD showed marked heterogeneity with height and throughout the day (Figure S6). Counterintuitively, WT *Ae. aegypti* unfed females demonstrated an increased probability to fly and rest at heights above 1m despite VPD increasing (i.e. increased desiccation pressure; GLM: *β* = 4.29, SE = 0.01, *z* = 288.0, p < 0.001; Figure 2C-D). Interestingly, and particularly for flight, *Ae. aegypti* showed a threshold-like relationship with VPD, with most mosquitoes moving above 1m at values above 1.5 kPa irrespective of the time of day (Figure 2C). VPD values ranging between 0.5 and 1.2kPa approximately correspond to indoor conditions (20 – 25 ºC; 40-60% RH) are generally considered comfortable for humans in which the air does not feel too dry or humid^73,74^. Values above 1.5kPa indicate very dry air and heating, which turn environments into less pleasant or risky for human health. Our interpretation of *Ae. aegypti* opposite behavioral response to VPD may be related to adults flying beyond their innate height range in search for better environmental conditions. Track analysis suggested this, given the higher frequency of flight downwards when VPD was higher than 1.5 (71% of tracks moved low) than when it was below that threshold (only 44 tracks moved low) (Figure S7). Analyses suggest *Ae. aegypti* females adjust their flight and resting height to minimize desiccation rather than to just relocate in areas that have lower temperature, as previously hypothesized.

**Figure 2.**
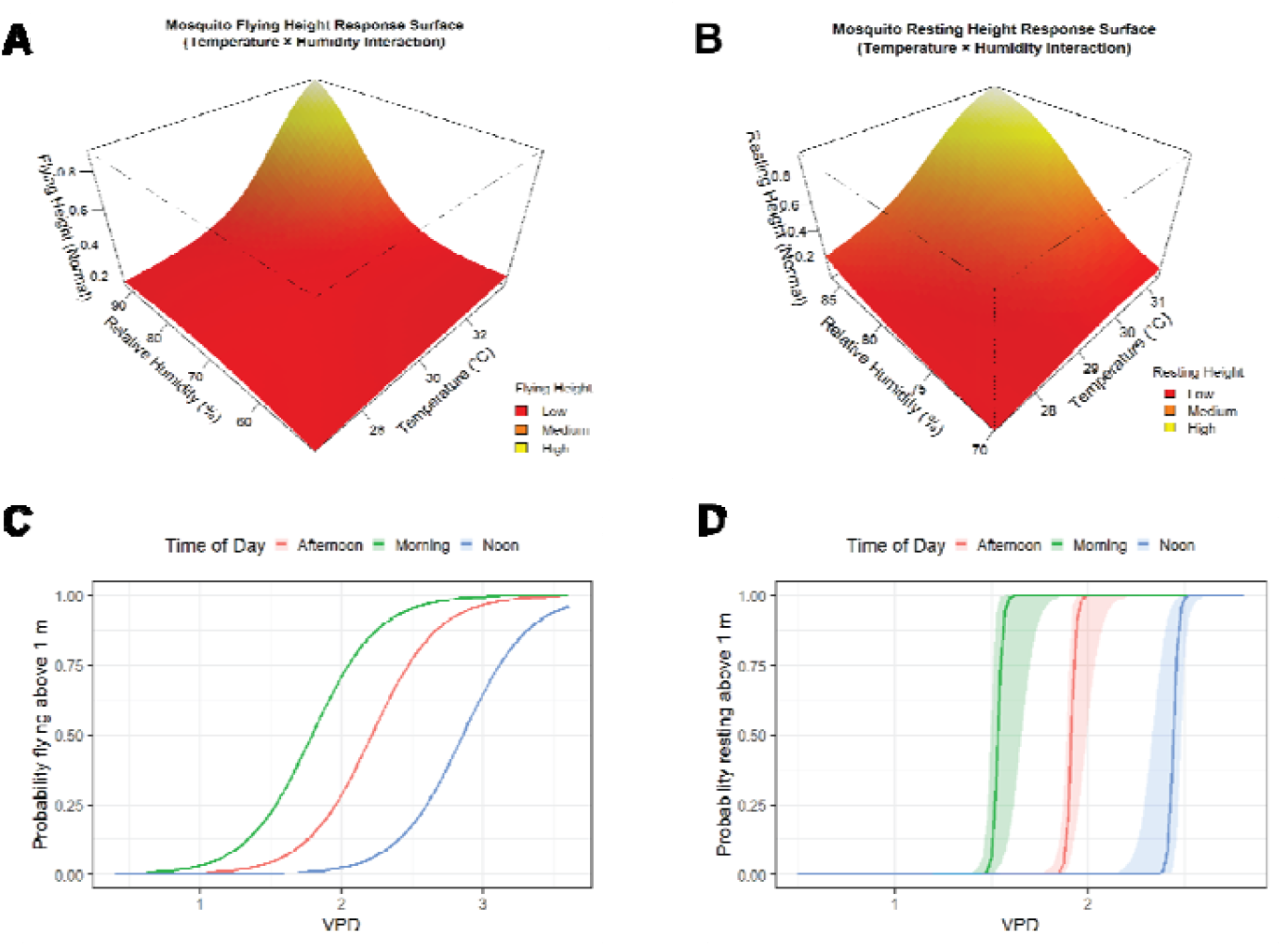
Interplay of microclimatic metrics (temperature, relative humidity, and VPD) and their impact on flying and resting height of Ae. aegypti mosquitoes (WT unfed females). **A** 3D GAM tensor surface plot visualizing the interaction between relative humidity (%) temperature (C), and their combined impact on flight height (track y-coordinates). Note flying height was normalized between 0 and 1 to allow use of the Beta error distribution. **B** 3D GAM tensor surface plot visualizing the interaction between relative humidity (%) temperature (C), and their combined impact on resting height. Like A, heights were normalized between 0 and 1 to allow use of the Beta error distribution. **C** Prediction plot of a Firth regression GLM demonstrating th relationship between increasing VPD (kPa) and the probability for mosquitoes (%) to rest above 1 meter, stratified by Time of Day (Afternoon replicates in red, Morning in green, and Noon in blue). **D** Prediction plot of a Firth regression GLM demonstrating the relationship between increasing VPD and the probability for mosquitoes to fly above 1 meter, stratified by Time of Day (Afternoon replicates in red, Morning in green, and Noon in blue).

### Color offsets *Aedes aegypti* innate preference for low resting heights

Although microclimatic conditions have been suggested to modulate flight and resting, these behaviors have not been considered within the context of other extrinsic conditions such as the color of resting sites. In experimental settings and the field, black targets are highly attractive to *Ae. aegypti* ^50,51^. We hypothesized that a behavioral shift in flight and resting may occur when colored targets are introduced at different heights, potentially offsetting other physiologic or innate responses. Figure 3 shows relative frequency of heights for resting and flying *Ae. aegypti* unfed females from WT and Lab strains, stratified by resting site color, which included homogeneous black and white as well as partial colors below or above 1m (see ‘Experimental Design’ in Methods). Consistent with the pattern described for unfed females exposed to white resting targets (Figure 1, Figure 3), when presented with black at low heights (i.e. below 1m), both flight and resting heights tended to occur primarily below 1 meter regardless of the mosquito strain (Figure 3, Table 2). However, when presented with black either 1) across all heights or 2) simply at high heights, (i.e. above 1m), WT unfed females demonstrated a significant shift in their flight heights above 1m (Black - *β* = 0.615 ± 0.07, *z* = 8.912, *p* < 2 x 10^-16^; Black:White - *β* = 0.159 ± 0.07, *z* = 2.296, *p* = 0.0217; Table S2, Figure 3). Although a similar impact was observed in resting height, it was only significant when unfed females were presented with black at high heights only (*β* = 1.61 ± 0.311, *z* = 5.183, *p* = 2.19 x 10^-07^, Figure 3, Table 2). Interestingly, most resting unfed females in the half-color treatments were caught right around the 1m mark, which is the line of contrast between both colors (Figure 3). Thus, it appears that highly attractive resting targets only have a significant impact on the resting preferences of mosquitoes rather than on their actual flight patterns, with such preference being even more pronounced if these highly attractive surfaces are presented at heights that oppose innate preferences and tolerable microclimatic conditions which we have shown are least favorable at heights above 1m (Figure S5).

**Table 1.**
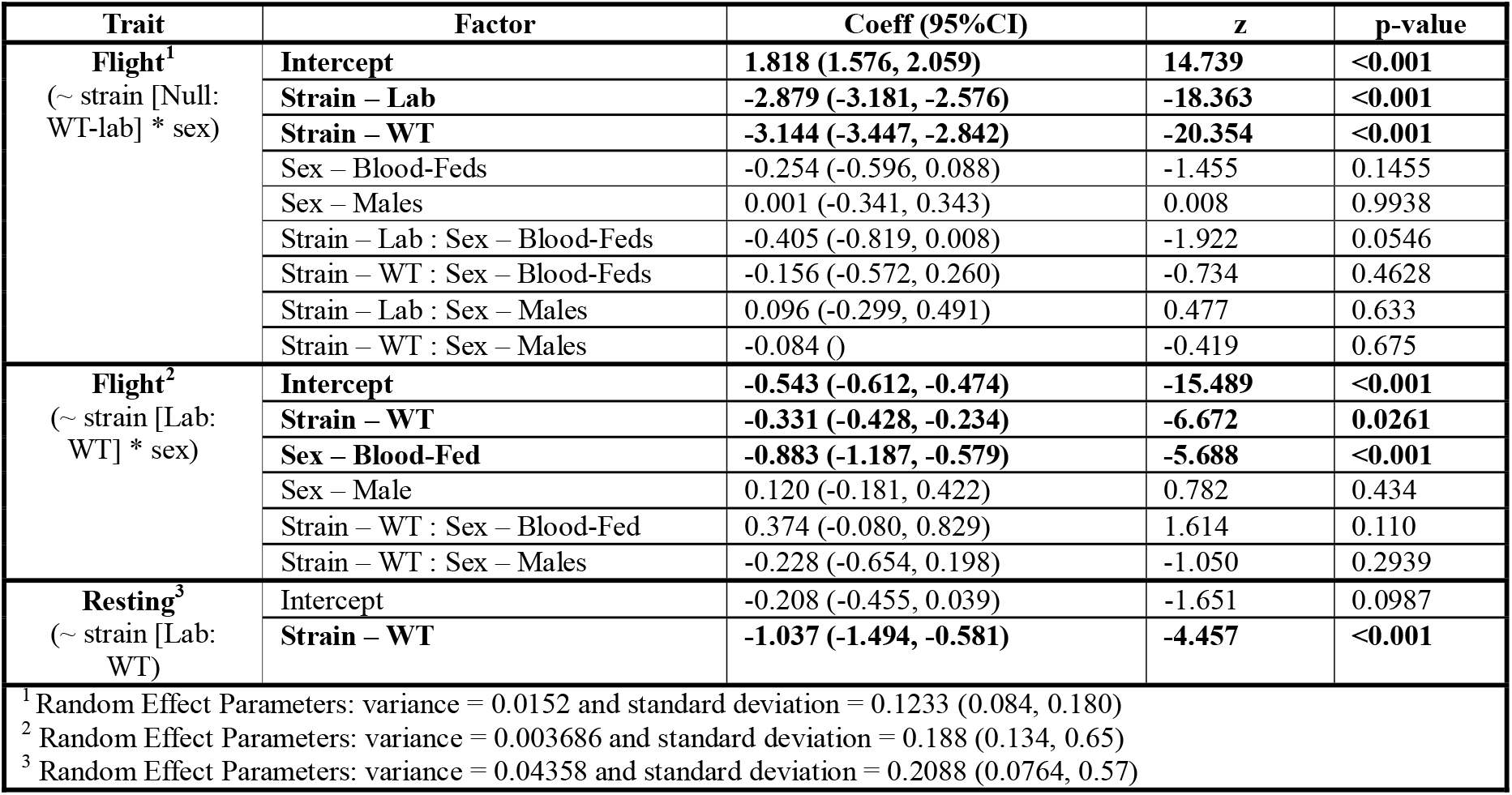
Drivers of *Aedes aegypti* endogenous innate flight and resting. Summarizing result outputs of the top performing linear mixed effects models fitting only endogenous factors (strain, sex/physiological status) for resting height and flying height. Models were fitted independently to each behavior and contained replicate as a random effect (1|replicate.id). Note that Unfed Females was set as reference for the fixed effect of Sex for both Flight and Resting. In the case of Strain, Null Simulated Random Walk was set as reference for the first Flight GLMM, while LAB was set as reference for the second Flight GLMM and the Resting GLMM.

| Trait | Factor | Coeff (95%CI) | z | p-value |
| --- | --- | --- | --- | --- |
| **Flight<sup>1</sup>**<br>(~ strain [Null: WT-lab] * sex) | Intercept | 1.818 (1.576, 2.059) | 14.739 | <0.001 |
|  | Strain – Lab | -2.879 (-3.181, -2.576) | -18.363 | <0.001 |
|  | Strain – WT | -3.144 (-3.447, -2.842) | -20.354 | <0.001 |
|  | Sex – Blood-Feds | -0.254 (-0.596, 0.088) | -1.455 | 0.1455 |
|  | Sex – Males | 0.001 (-0.341, 0.343) | 0.008 | 0.9938 |
|  | Strain – Lab : Sex – Blood-Feds | -0.405 (-0.819, 0.008) | -1.922 | 0.0546 |
|  | Strain – WT : Sex – Blood-Feds | -0.156 (-0.572, 0.260) | -0.734 | 0.4628 |
|  | Strain – Lab : Sex – Males | 0.096 (-0.299, 0.491) | 0.477 | 0.633 |
|  | Strain – WT : Sex – Males | -0.084 () | -0.419 | 0.675 |
| **Flight<sup>2</sup>**<br>(~ strain [Lab: WT] * sex) | Intercept | -0.543 (-0.612, -0.474) | -15.489 | <0.001 |
|  | Strain – WT | -0.331 (-0.428, -0.234) | -6.672 | 0.0261 |
|  | Sex – Blood-Fed | -0.883 (-1.187, -0.579) | -5.688 | <0.001 |
|  | Sex – Male | 0.120 (-0.181, 0.422) | 0.782 | 0.434 |
|  | Strain – WT : Sex – Blood-Fed | 0.374 (-0.080, 0.829) | 1.614 | 0.110 |
|  | Strain – WT : Sex – Males | -0.228 (-0.654, 0.198) | -1.050 | 0.2939 |
| **Resting<sup>3</sup>**<br>(~ strain [Lab: WT]) | Intercept | -0.208 (-0.455, 0.039) | -1.651 | 0.0987 |
|  | Strain – WT | -1.037 (-1.494, -0.581) | -4.457 | <0.001 |
| <sup>1</sup> Random Effect Parameters: variance = 0.0152 and standard deviation = 0.1233 (0.084, 0.180) |  |  |  |  |
| <sup>2</sup> Random Effect Parameters: variance = 0.003686 and standard deviation = 0.188 (0.134, 0.65) |  |  |  |  |
| <sup>3</sup> Random Effect Parameters: variance = 0.04358 and standard deviation = 0.2088 (0.0764, 0.57) |  |  |  |  |

**Table 2.**
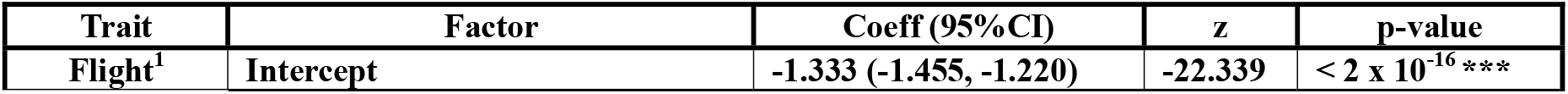
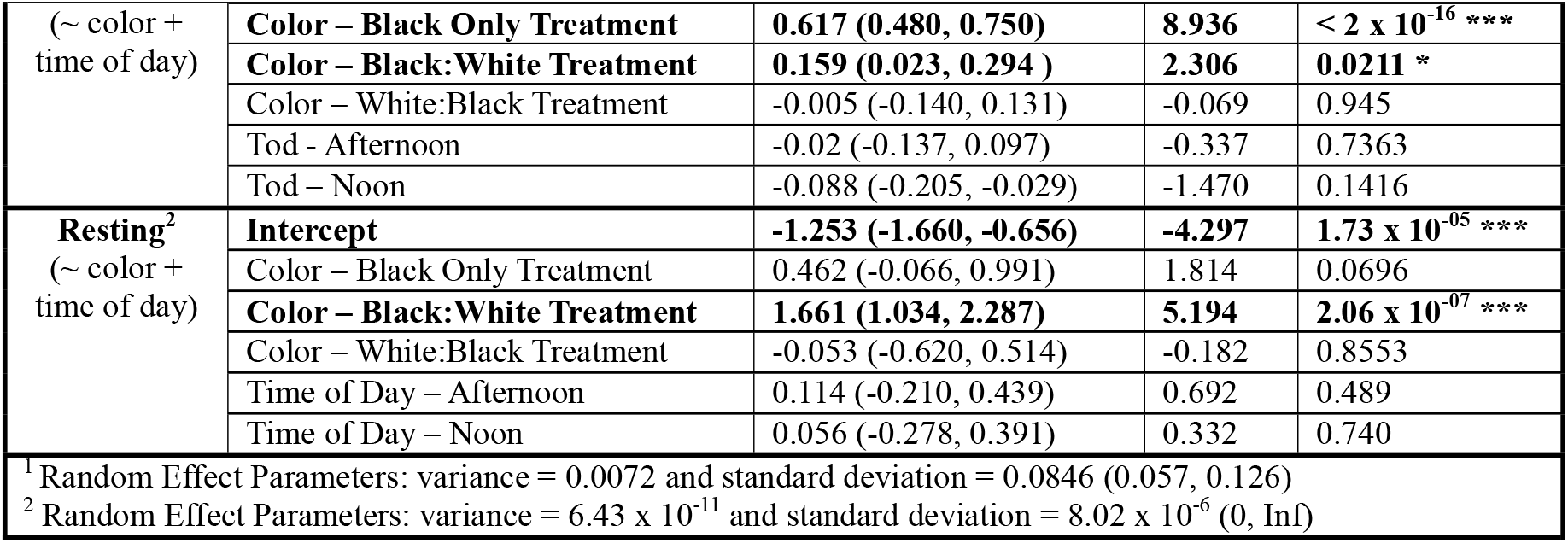
Exogenous drivers of *Aedes aegypti* flight and resting behavior. Summarizing results of the top performing linear mixed effects models fitting only exogenous factors (color and time of day) for resting height and flying height. Models were fitted independently to each behavior and contained replicate as a random effect (1|replicate.id). Note that for both flight and resting models, “White” was set as reference for the fixed effect of color treatment, and “Noon” as the reference for the fixed effect of time of day (proxy for microclimate).

| Trait | Factor | Coeff (95%CI) | z | p-value |
| --- | --- | --- | --- | --- |
| Flight<sup>1</sup> | Intercept | -1.333 (-1.455, -1.220) | -22.339 | < 2 x 10<sup>-16</sup> *** |
| (~ color +<br>time of day) | **Color – Black Only Treatment** | **0.617 (0.480, 0.750)** | **8.936** | **< 2 x 10<sup>-16</sup> \*\*\*** |
|  | **Color – Black:White Treatment** | **0.159 (0.023, 0.294 )** | **2.306** | **0.0211 \*** |
|  | Color – White:Black Treatment | -0.005 (-0.140, 0.131) | -0.069 | 0.945 |
|  | Tod - Afternoon | -0.02 (-0.137, 0.097) | -0.337 | 0.7363 |
|  | Tod – Noon | -0.088 (-0.205, -0.029) | -1.470 | 0.1416 |
| **Resting<sup>2</sup>**<br>(~ color +<br>time of day) | **Intercept** | **-1.253 (-1.660, -0.656)** | **-4.297** | **1.73 x 10<sup>-05</sup> \*\*\*** |
|  | Color – Black Only Treatment | 0.462 (-0.066, 0.991) | 1.814 | 0.0696 |
|  | **Color – Black:White Treatment** | **1.661 (1.034, 2.287)** | **5.194** | **2.06 x 10<sup>-07</sup> \*\*\*** |
|  | Color – White:Black Treatment | -0.053 (-0.620, 0.514) | -0.182 | 0.8553 |
|  | Time of Day – Afternoon | 0.114 (-0.210, 0.439) | 0.692 | 0.489 |
|  | Time of Day – Noon | 0.056 (-0.278, 0.391) | 0.332 | 0.740 |
| <sup>1</sup> Random Effect Parameters: variance = 0.0072 and standard deviation = 0.0846 (0.057, 0.126) |  |  |  |  |
| <sup>2</sup> Random Effect Parameters: variance = 6.43 x 10<sup>-11</sup> and standard deviation = 8.02 x 10<sup>-6</sup> (0, Inf) |  |  |  |  |

**Figure 3.**
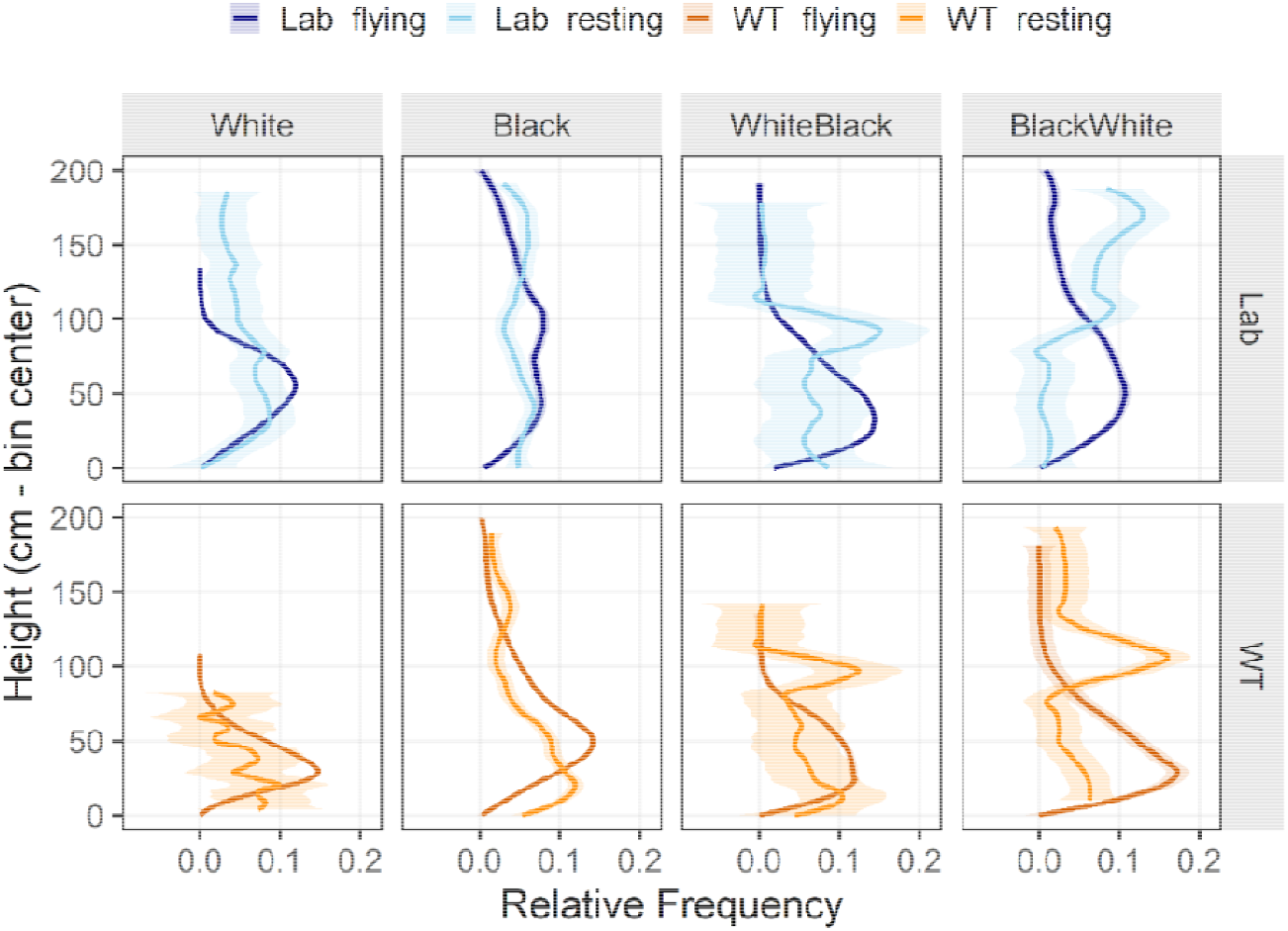
Influence of resting target color on *Ae. aegypti* flight and resting height preferences stratified by strain. LOESS-smoothed curves show the relative frequency of flying and resting across vertical height bins. Each panel represents a combination of strain (Lab vs WT) and color treatment. For Lab, flying is represented in dark blue while resting is represented in light blue. In the case of WT, flying is represented in dark orange, while resting is represented in light orange. The y-axis indicate height (bin center in cm) and the x-axis indicate the relative frequency of observations within that height range. Shaded ribbons represent 95% confidence intervals around the fitted LOESS curves.

**Figure 4.**
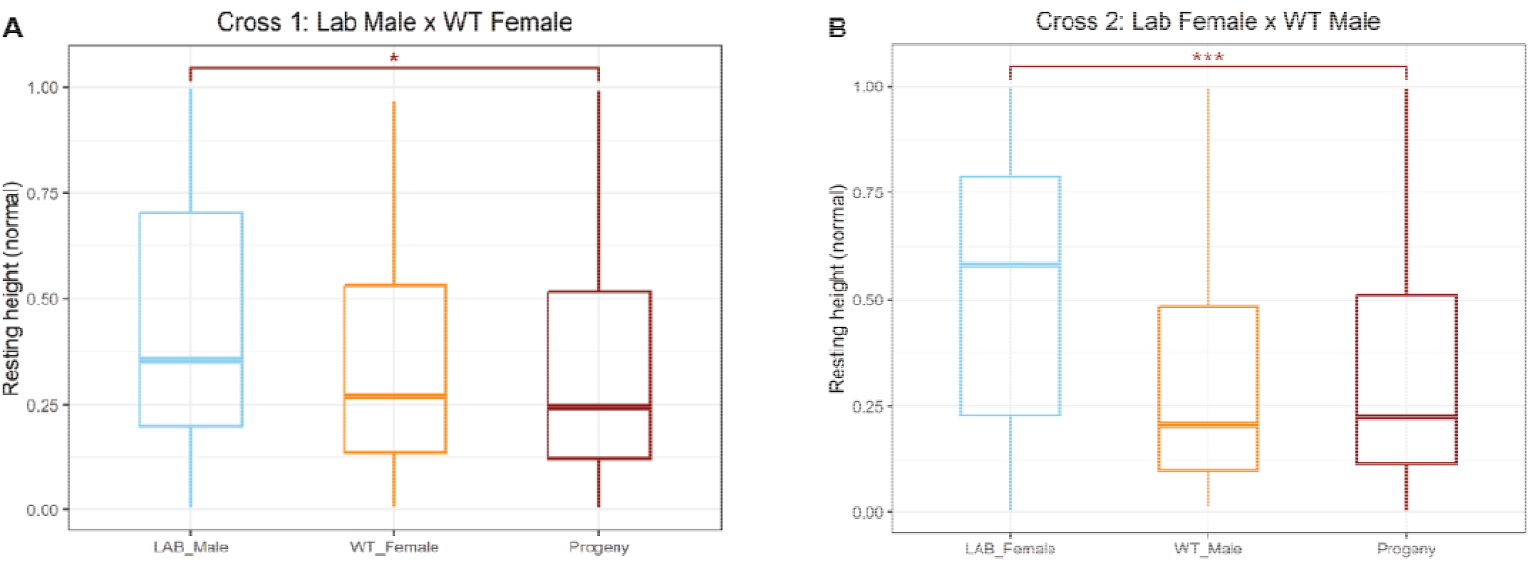
Boxplots summarizing resting height measurements between reciprocal cross progeny and their respective parents. **A** presents results from first crossing between Lab Males (skyblue) and a WT Females (orange) and their progeny (both males and females; red). **B** demonstrates results form the second crossing between Lab Females (skyblue) and WT Males (orange) and their progeny (both males and females; red).

### Patterns between sexes and physiological states

Considering endogenous factors only, when males of either strain (Lab-adapted vs WT) where presented with non-attractive sticky resting surfaces (uniform White), they flew and rested quite similar to the unfed females (fly: *β* = -0.12 ± 0.154, *z* = 0.78, *p* = 0.434, rest: *β* = -0.172 ± 0.21, *z* = -0.822, *p* = 0.411 Table 2; Table S1; Figure S8). Although this was also the case for WT blood-fed females (fly: *β* = 0.374 ± 0.23, *z* = 1.614, *p* = 0.107; rest: *β* = 0.131 ± 0.83, *z* = 0.158, *p* = 0.875; Table S4; Figure S8), the flying height of Lab-adapted blood-fed females was significantly lower than Lab unfed females (fly: *β* = -0.883 ± 0.155, *z* = -5.69, *p* = 1.28 x 10^-8^; Table S4; Figure S8), demonstrating a significant interaction between a strain’s genetic background and the physiological demands of digesting a full blood meal only on flight height. As we know, the genetic background of the Lab strain we used in the study (Rockefeller) is highly impacted by the bottleneck effects of continuous colony rearing resulting in decreased genetic variation ^62,63^. This behavioral difference may be directly influenced by the continuous artificial selection and blood-feeding of mosquitoes under controlled Lab conditions.

When presented with attractive sticky resting surfaces (i.e. uniform Black, or either half-black condition) no significant interaction between strain (Lab) and physiological status (Blood-Fed) impacting flight and resting height is observed (flight: *β* = -0.0163 ± 0.15, *z* = -0.108, *p* = 0.9137). Instead, similar to the flight results of unfed females shown in Figure 3, both males and blood-fed females (regardless of strain), demonstrated a significant increase in flight height, but only when Black was present above 1m (Uniform Black - flight: *β* = 0.820 ± 0.124, *z* = 6.60, *p* = 3.87 x 10^-11^, rest: ; Black:White – flight: *β* = 0.496 ± 0.124, *z* = 3.998, *p* = 6.39 x 10^-5^). Although sex/physiological status was not a significant fixed effect driving resting height preferences, there was a slight difference in the significant effect of color treatment. Despite a similar impact of Black above 1m (Black:White) increasing resting height (*β* = 1.055 ± 0.1388, *z* = 7.645, *p* = 2.10 x 10^14^), when uniformly presented with Black, mosquitoes still preferred to rest low just like they would in the absence of any attracting surfaces (*β* = 0.219 ± 0.134, *z* = -1.639, *p* = 0.1012). Moreover, these results once again indicate a significant reduction in observed resting height when mosquitoes (regardless of strain and or sex/physiological status) are presented with Black just below 1m (White:Black; *β* = -0.304 ± 0.136, *z* = -2.24, *p* = 0.025). Taken together, our results record how resting targets can modulate mosquito flight under semi-field conditions, and point to a highly conserved affinity to black surfaces regardless of genetic background. Moreover, these results indicate adjacently contrasting color surfaces have a higher impact on the resting height rather than the flight of *Aedes aegypti* mosquitoes, while considering sex/physiological status and strain. In line with our microclimatic and VPD results from above, time of day did not seem to have a significant effect when the models also consider color and the endogenous factors (strain and sex/physiological state), with the only exception of “Noon” replicates only having a significant reduction in flight height (*β* = -0.219 ± 0.106, *z* = -2.057, *p* = 0.0397). This suggests mosquitoes may be responding to rising temperatures (and in turn, decreasing relative humidities) which typically reach a daily peak around midday, thus becoming less interested in flying at higher heights (above 1 meter).

### Progeny from crosses strongly resemble WT *Aedes aegypti* resting behavior

Given the consistency for the Lab mosquitoes to fly and rest significantly higher than the WT mosquitoes, even while accounting for exogenous factors, we produced two reciprocal crosses using both strains as an initial step in investigating possible genetic/epigenetic factors driving and/or modulating mosquito resting and flight behavior (see ‘Experimental Design’ in Methods). Progeny from the first crossing rested significantly lower (*β* = -0.606 ± 0.119, *z* = -5.09, *p* = 3.51 x 10^-07^) in comparison to their Lab parent (Male - *β* = 0.394 ± 0.181, *z* = 2.176, *p* = 0.0296) but not from the WT parent (Female - *β* = -0.0001 ± 0.197, *z* = -0.001, *p* = 0.999; Table S6). This significant difference was even more pronounced in the second crossing, with the progeny resting height much lower (*β* = -0.494 ± 0.098, *z* = -5.038, *p* = 4.7 x 10^-07^) than their Lab parent (Female; *β* = 0.516 ± 0.146, *z* = 3.546, *p* = 4 x 10^-4^; Table S6). The recapitulation of the WT behavior after crosses may be indicative of potential heritable marker(s) from a WT genetic background modulating behavior. While more research will be needed to ascertain whether epigenetic signaling influences *Ae. aegypti* flight and resting, our findings provide important insight about the loss of traits in highly inbred mosquito colonies. Findings from our study provide additional justification for the back-crossing of laboratory mosquito strains containing *Wolbachia* or transgenic genes with WT strains, given their flight and resting behavior may be significantly impaired when inbred laboratory strains are used.

### Study Limitations

We acknowledge several limitations of our study. We did not have any control over wind and lighting conditions within the hut, which are both known to impact mosquito movement ^75–77^. In addition, we recognize the height limitations of the hut (i.e. 2 meter) may not be an all-encompassing representation of the height structure of most homes in Merida. Moreover, the PFMD device we used to record the mosquitoes contains a significant blind spot, which did not allow us to comprehensively record the mosquitoes across the entire space inside of the hut. Field technicians validated 2.5 meters from the front wall of the hut as the minimum distance for an object to be in complete view of both PFMD cameras. While one of the major findings of our study is the strong strain difference in behaviors, we only tested for one strain effect (a Merida-derived mosquito population). Future studies focusing on mosquito strains from different settings and contexts (i.e., dry versus humid environmental conditions) may provide better insights into the behavioral fine-tuning that occur as *Ae. aegypti* adapts to different conditions.

## Conclusions

While particular flight and resting preferences of *Ae. aegypti* have been proposed as consequence of microclimatic preferences ^45,67^, empirical field evidence remains limited. We demonstrate that wild and laboratory *Ae. aegypti* strains differ markedly in flight and resting height preferences. These differences persist under similar environmental conditions and are partially recovered in reciprocal crosses, revealing a likely heritable component of behavioral divergence. Furthermore, by quantifying microclimatic gradients and VPD, we show that mosquitoes dynamically adjust their vertical position to minimize desiccation risk, while strong visual cues (black surfaces) can override these microclimatic constraints. Together, these results establish a mechanistic framework that links genetics, physiology, and environment to explain *Ae. aegypti* spatial behavior indoors.

Our findings have important relevance for the design of novel vector control interventions and for the evaluation of existing ones. The signal of a GxE interaction in flight and resting suggest that such behaviors may be heritable. Behavioral resistance to vector control has been documented for *Anopheles* spp in sub-Saharan Africa^78^, particularly as changes in feeding pattern (increased exophily of *Anopheles arabiensis* leading to failure of indoor residual spraying^79^) or shifts in diel towards earlier biting (*Anopheles farauti* in the Solomon Islands shifting feeding towards earlier times^80^. While several other examples of behavioral response to interventions have been documented^78^, evidence of potentially heritable behaviors has lacked from such assessments. Our findings suggest that interventions that selectively apply insecticides indoors, such as Targeted Indoor Residual Spraying^81^ may inadvertently select for high resting, provided there is a fitness advantage for surviving adults resting beyond the area where insecticides are applied. As such, monitoring for indoor flight height may be a requisite for any intervention that is selectively deployed indoors. Other important insight from this study that can help inform interventions is the preference of Ae. aegypti for specific resting cues, such as contrasting colors. Mosquitoes showed changes in innate free-flight in the presence of such “targets”, providing potential for the development of novel approaches for targeting resting that do not imply insecticide spraying.

## Methods

### Experimental setup

An experimental hut located in the campus of the Autonomous University of Yucatan, Mexico, measuring 3×4×2m (WidthxLengthxHeight) was fitted with protective screens to prevent mosquitoes from entering or escaping (SI Text and Figure S1). Microclimatic conditions within the hut were recorded at four different heights using HOBO Temperature and Relative Humidity (UX100-003, Onset Computer Corporation, United States) loggers. Loggers were strapped to the West wall of the hut at 0m, 0.64m, 1.28m, and 1.92m. Each HOBO was configured to take a temperature and relative humidity measurement every ten minutes. *Aedes aegypti* free flight within the hut was mapped using the Photonic Fence Monitoring Device, PFMD, a device capable of detecting and tracking insects ranging in size from 4 mm to 40mm with high accuracy (Patt et al., 2024). Thanks to a pair of stereoscopic infrared cameras, which allow identifying subjects from silhouettes generated by the reflection of mosquitoes against a retroreflector screen, the PFMD can detect and map in 3D the position of thousands of mosquitoes at a frame detection frequency of 1/100 of a second. The SI Text provides a detailed description of the PFMD architecture and details of the algorithms for mosquito identification and track assembly. Mosquito resting height was measured by installing four sticky traps measuring 30m wide and 2m tall and consisting of white plastic tablecloth covered with Catch EM’ Sticky Insect Trap Coating (Bug Ball, Inc, United States), placed at each corner of the hut (Figure S2) to sample resting mosquitoes concurrently to the PFMD recording their flight activity. White, black and white-black sticky traps were used depending on the experimental treatment. Traps were replaced every week according to the corresponding color condition for that particular week (see SI Text for more details).

### Experimental Design

A factorial experimental design included endogenous (mosquito genetic background, sex, physiological state) and extrinsic (color of resting sites, temperature, humidity) factors we hypothesized are determinants of *Ae. aegypti* free-flight and resting height preference. Three replicates of each condition were performed, and values across replicates were used for analysis. The study initially quantified the free flight and resting behaviors of *Ae. aegypti* under no external stimuli (white color resting) as a way of quantifying innate behaviors. Mosquito genetic background had two levels, a laboratory strain (Rockefeller from CDC, termed as Lab) and a field-derived strain (termed as wild-type, WT). Details of each strain and rearing conditions are in SI Text. Afte characterizing *Ae. aegypti* innate behavior, secondary experiments quantified the impact of extrinsic factors such as microclimate (temperature and humidity) as well as the presence of visual cues that may influence flight and resting (the presence of black surfaces). All experimental conditions were repeated for unfed females, males, and bloodfed females.

The experimental design involved the release of 100 mosquitoes of each specific strain, sex (Males – M vs Unfed Females – UF) or physiological state (UF vs Blood-Fed Females – BF), deprived from sucrose for at least 18 hours, within the center of the hut. To ensure BF obtained a complete blood-meal, all 100 individuals were fed by arm two-hours prior to being taken into the experimental hut. Once brought to the hut, mosquitoes are released (using a manual remote mechanism from the outside of the hut) to start the experiment and recording with the PFMD. Monitoring occurred for a period of 2 hours, at which time the PFMD was turned off and one field technician entered the hut with a Prokopack aspirator^49^ to collect all free flying mosquitoes from the hut and retrieve all mosquitoes adhered to the sticky traps, registering the height of each individual mosquito with a measuring tape. Given that *A. aegypti* has been shown to follow a bimodal crepuscular activity pattern (i.e. peak activity occurs at dawn and then at dusk^82^) replicates were scheduled in a manner allowing us to capture variation in behavior due to this diel activity pattern (SI Text). A grand total of 72 replicates were carried out, involving 36 groups of 100 individuals (12 for each experimental group – M, UF, and BF). Details of the procedures for releasing mosquitoes, monitoring them within the hut and processing them after the experiment ended are described in SI Text.

### Modeling mosquito random flight

One of the challenges of analyzing behavioral data is that it is hard to compare observed patterns to expected behaviors. For mosquito flight, we generated a null model of flight based on a 3D random walk process^83^. The output of the PFMD (consisting of 3D coordinates for each mosquito position at 100 Hz) was processed into vector component distances at each discrete step along three spatial axes in centimeters (i.e. the change in x, y, and z from location to location: dx, dy, dz). From this processed data we further calculated 3D step-lengths (SL) associated with each successive relocation following:

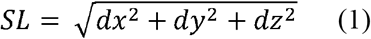

Stepwise turning angles were also calculated in 3D space. At each step we determined the absolute turning angles, and subsequently the relative turning angles^84^, along both the XY plane and the XZ plane, resulting in two relative turning angle measures per movement (i.e. relative turning angles associated with both ‘pitch’ and ‘yaw’). This multi-planar methodological approach was partially motivated by the expectation that gravity and/or other factors might result in differing turning angle distributions with respect to these planes of movement.

Empirical step-length data was then used to parameterize a theoretical gamma distribution of the form:

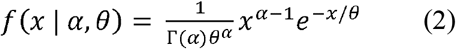

where α is the shape parameter, θ is the scale parameter and Γ the gamma function. Two separate theoretical turning angle distributions following a Von Mises distribution were parameterized by empirical data along each respective axial plane, with one XY distribution and one XZ distribution. The Von Mises distribution took the form:

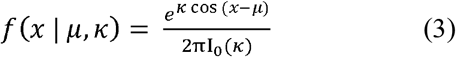

where *μ* is the center of the distribution, *κ* is a concentration parameter, and I_0_(*κ*) is the modified Bessel function.

Using these empirically informed theoretical distributions, we simulated mosquito movement based on a 3D random-walk replicating multi-planar turning angle behaviors and step-length displacements at the sub-second temporal scale. More details of model parameters, code, and outputs are found in the SI Text.

### Statistical Analyses

Different analyses were conducted for flight versus resting data, and a detailed description of all statistical tests and their rationale is provided in SI Text.

Generalized linear mixed models (GLMMs) were used to further assess the individual and joint effects of strain, experimental group, color treatment, and time of day over a) resting height, and b) flight track height (y) coordinates. All models were fitted using a Beta error distribution with a logit link function, and “replicate id” as the only random effect. To further investigate the collinearity of temperature and relative humidity and their combined effect on resting height, we fitted Generalized Additive Models (GAMs). Four kinds of models were built – 1) mean temperature and mean relative humidity as a tensor; 2) mean temperature as a smoothing term; 3) mean relative humidity as a smoothing term, and 4) estimated vapor pressure deficit (VPD) as a smoothing term. Vapor pressure deficit (VPD) considers both temperature and relative humidity and quantifies the drying power of the air. VPD is a biologically relevant measure of desiccation stress experienced by mosquitoes^65^. Details for calculating VPD are found in SI Text. All of these models were fitted with the Beta distribution as the link function and set to use the Restricted Maximum Likelihood (REML) method for parameter estimation.

## Ethics Statement

No human or vertebrate subjects of any kind were involved in the study presented here. The authors manifest no conflicts of interest regarding the presented research.

## Data accessibility

All datasets and R scripts used for statistical analyses and figure generation in this study are available in an open-access GitHub repository: https://github.com/davidjim890/Drives-of-mosquito-indoor-free-flight-and-resting.git.

## Notes

https://zenodo.org/records/17817358

### Competing Interest Statement

The authors have declared no competing interest.

### Funding

- National Institute of Allergy and Infectious Diseases: U01AI148069, T32AI138952

