## Supplementary materials for "Drivers of mosquito free-flight and resting behavior indoors"

**Authors and Affiliations**

Not listed for double anonymous peer-review (DAPR)

### Supplementary Methods

#### Experimental Setup

The UCBE (Unidad Colaborativa de Bioensayos Entomológicos) is an independent academic and research center situated within Autonomous University of Yucatan’s Campus of Biological and Agricultural Sciences in Mérida, Yucatán, Mexico. Established in 2014, UCBE plays a pivotal role in evaluating and advancing vector surveillance and control strategies, with a strong focus on Aedes aegypti—the primary vector for dengue, chikungunya, and Zika. The unit is officially recognized by Mexico’s Ministry of Health as a reference center and is recognized as a WHO GLP (Good Laboratory Practices) Evaluation Center.

The experimental huts (Figure S1) measure 4m x 2m x 3m and its walls are made of PVC panels (thickness 2cm) affixed to an aluminum frame. The entire hut system is housed under a PVC roof, which prevents rainfall and debris from entering the hut. Huts are located within 100m of the UCBE laboratory (Figure S1), within a small patch of dry forest and exposed to the climate typical of the Yucatan Peninsula. A tarp was placed on one end of the hut to protect the computers used to track mosquito behaviors from direct sun exposure and rainfall.

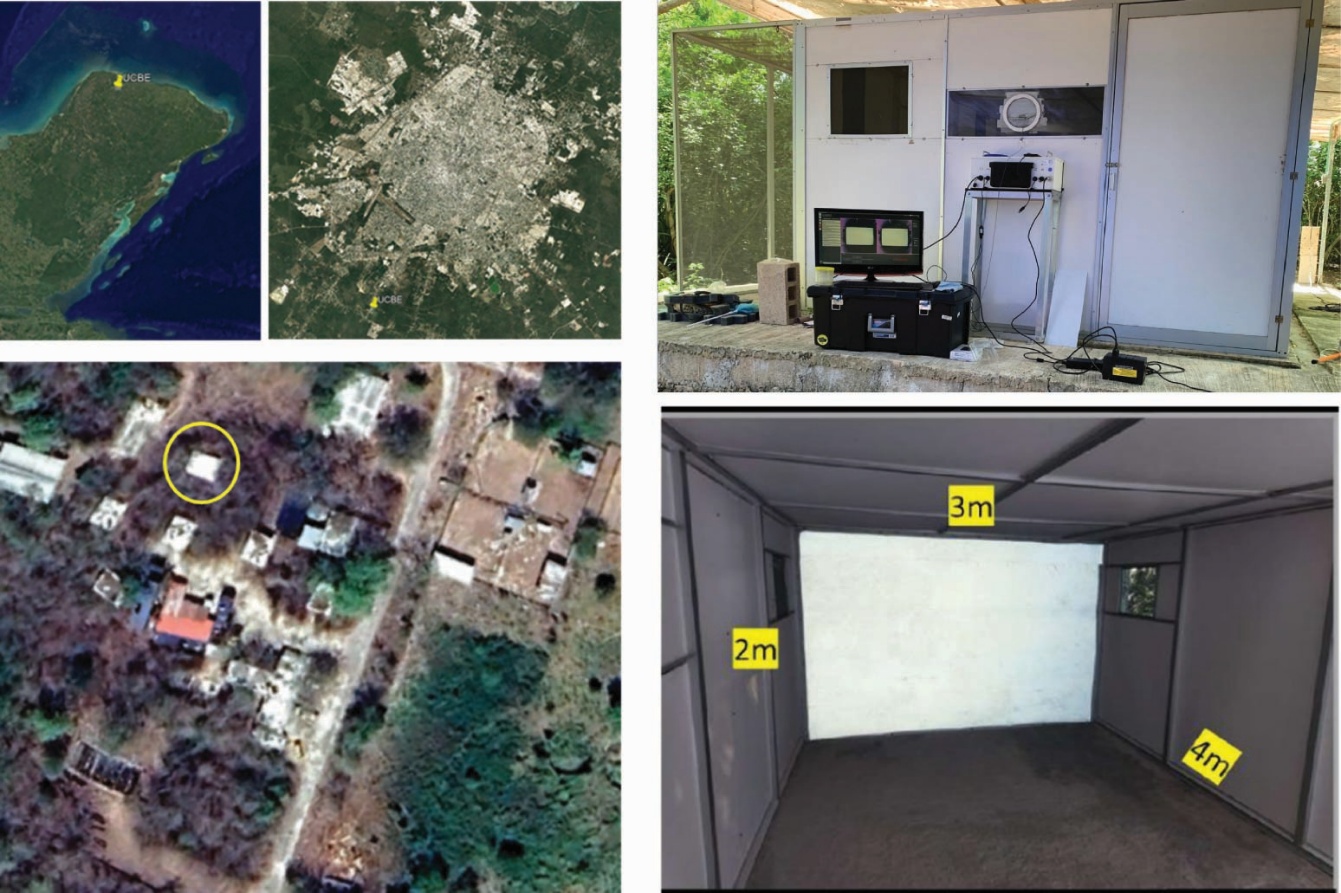
 **Figure S1. Location of experimental huts within UCBE (yellow circle on low-left) and of UCBE within Merida and Mexico (top-left). External arrangement of experimental hut (top-right) and inside view showing the placement of the reflective fabric (bottom-right).**

Within the hut, a sticky paper was used to measure resting height. Four 20cm x 2m strips of paper were placed in the hut (one on each corner) and coated with Catch EM’ Sticky Insect Trap Coating (Bug Ball, Inc, United States) (Figure S2). Such layout was successfully used in Thailand to track mosquito resting indoors^1^, and has shown no apparent behavioral impact on mosquito behavior in such study and our evaluation.

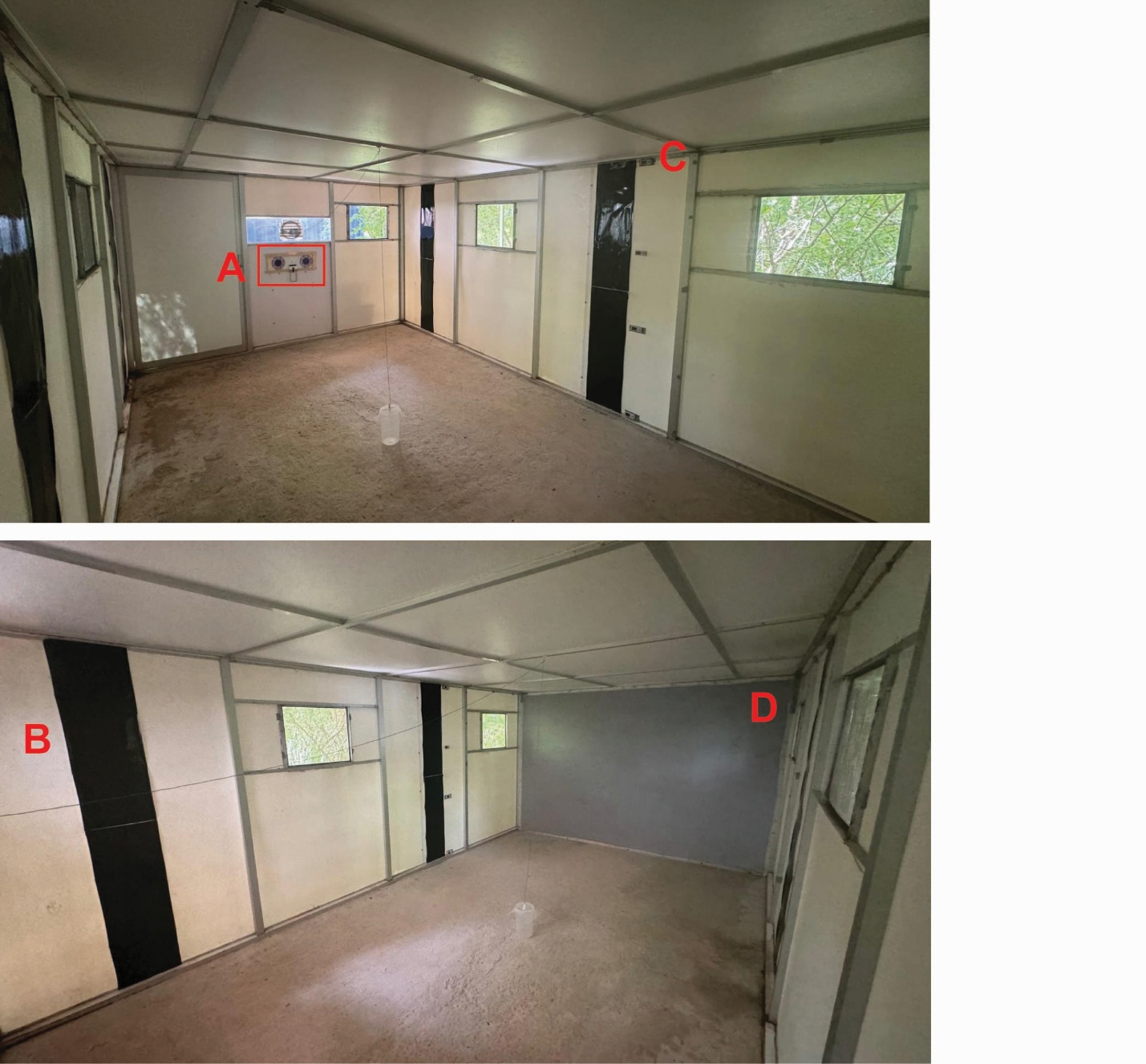

**Figure S2. Layout of the experimental hut, indicating the placement of the PFMD stereoscopic mapping tool (A), the sticky traps (B), the HOBO loggers (C) and the reflective fabric (D).**

#### Stereoscopic mosquito tracking with PFMD

The company Photonic Sentry, with support from Global Health Labs and the Bill &

Melinda Gates Foundation, has recently developed a device able to track insect flight,

enabling more detailed study of mosquito behavior from a version of a tracking

system previously described^2^. The Photonic Fence Monitoring Device (PFMD)

is capable of detecting and tracking insects such as mosquitoes, bees, and flies,

ranging in size from approximately 4 mm to 40mm. The device’s pair of offset

stereoscopic cameras identify the three-dimensional location of a target (Fig S3A). Each

stereoscopic camera is equipped with a circular arrangement of infrared LEDs, which

allows identification of subjects from the size of their silhouettes generated by the

infrared LED reflection against a retroreflector screen. This onboard IR illumination not

only allows the device to be used during the day, but also during the night. The data can

be collected on a continuous 24/7 basis for days at a time (up to 7 days or longer,

depending on data density) and then visualized and analyzed through Photonic Sentry’s

own cloud-based, online researcher portal. If desired, the data can also be exported in

comma-separated value (CSV) format and imported into software like R for other kinds

of analysis.

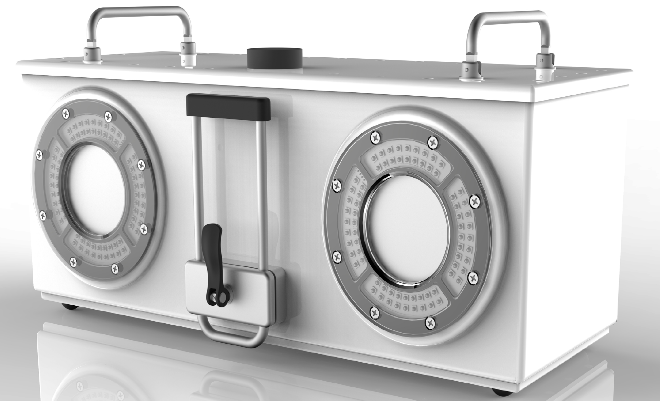

**Figure S3A.** Frontal image of the Photonic Fence Monitoring Device unit

The active region of the device involves a truncated square pyramid that is 3m x 3m x RANGE, where RANGE is set at the time of factory calibration varying from 3m to 10m (Fig S3B). Depending on which RANGE is used (3m vs 10m), the device will experience a blind spot in which subjects will not be detected. In the case that RANGE is 3m, the device will not be able to detect any insects at a range less than 1m. If the range was set to 10m then subjects that are at a range less than 5m will not be detected by the PFMD. At the moment of recording, the PFMD will use a reference point (in between both cameras) to set the z-axis, determine the distance to the retroflector screen, and set the active region. This is important to note because although angling the device will not alter the shape of the pyramid, it will alter the orientation of the z-axis, and thus, the field of view.

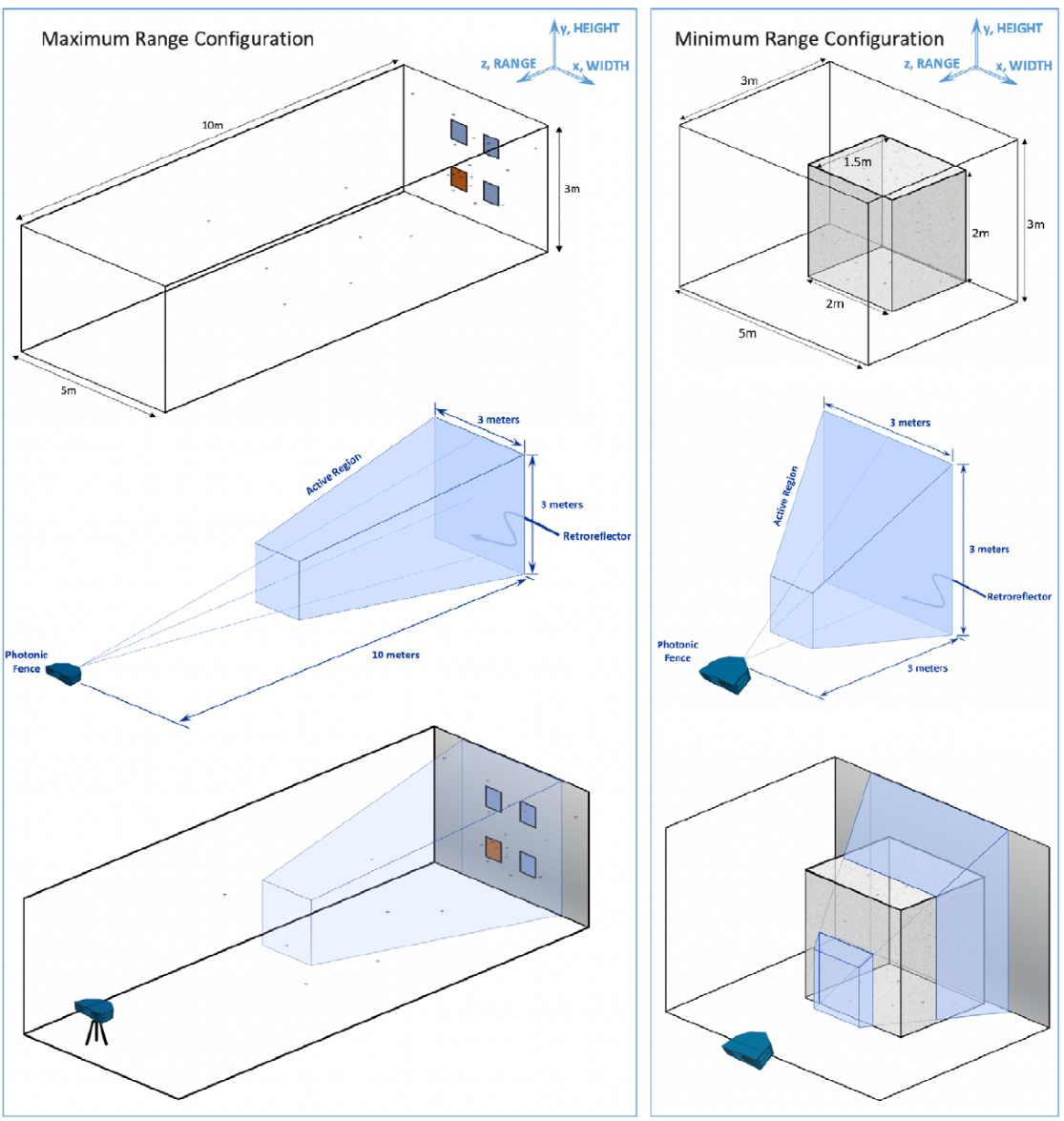

**Figure S3B.** A graphical representation of the square pyramid used by the Photonic Fence Monitoring Device to establish its active recording region at both minimum (3m) and maximum (10m) range configurations.

The device can be set to record either indefinitely (must be manually stopped) or for a

predetermined period of time. Once the device is recording, it is preferable that no

obstructions enter the active field in order to avoid noise in the data. Once the data has

been gathered and uploaded to the cloud-server, there are multiple ways of visualizing

what the PFMD recorded. The first of these is a 3D visualization of flight tracks

displayed as lines over time with a filled circle representing the origin of the flight and an

X representing the termination (Fig S3C). Track observation time, speed, acceleration,

or size can be all be viewed using the “Colorize Tracks By” menu within the portal. In

addition to this, the portal will also generate a 3D heatmap displaying the density

distribution of track observations, showing the relative likelihood of observing an insect

in a given position (Fig S3D). 3D heatmaps can also be observed for speed and

acceleration, which will show the average of each property interpolated across the

volume. Lastly, the portal will generate histograms showing track-level aggregate

including insect size, track lengths, track durations, track speed, and track acceleration

across all observations (Fig S3E).

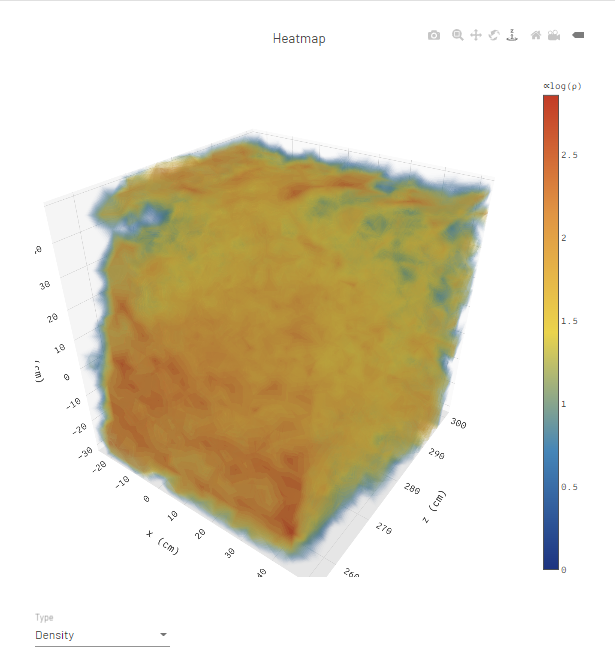

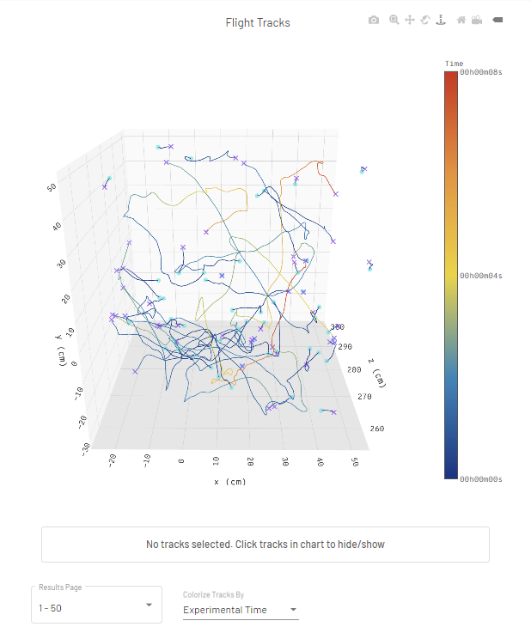

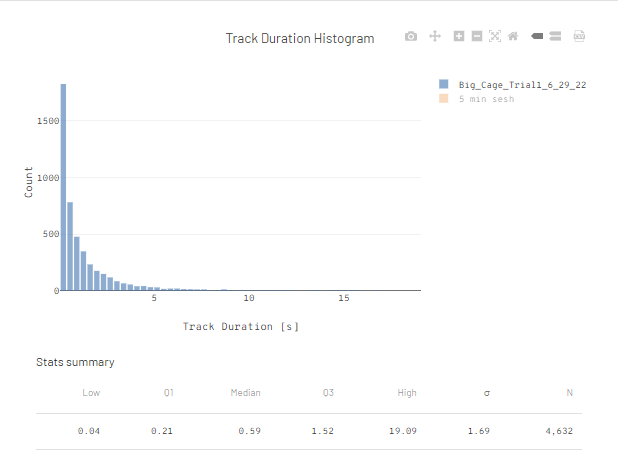

**Figure S3C-E. Top left image (S3C) is an example of the Flight Track graph generated by the**

**researcher portal visualizing 3D paths for tracks recorded during that particular trial. Top right**

**image (S3D) demonstrates the generated heatmap from all of the tracks in S3C with blue**

**representing volume where subjects are least likely to be observed and red where they are**

**most likely to be observed. Bottom image (S3E) is an example of the Track Duration Histogram**

**generated by the researcher portal.**

#### Mosquito strains

As part of the study, we evaluated two strains of *Aedes aegypti* coming from very distinct genetic backgrounds. A laboratory-adapted *A. aegypti* strain obtained from collaborators at the Center for Disease Control and Prevention (CDC) (Rockefeller – *ROCK*) of which a colony has been established and maintained at UCBE. Given that this strain has experienced strong selection to withstand and thrive in standardized laboratory conditions, it is likely that traits driving resting and flight height preferences have not been the primary target of such selection. In other words, given that resting and/or flight height preferences do not directly impact the survival or reproductive output of ROCK mosquitoes in laboratory conditions, selection is unlikely to act and may even remove particular traits driving such preferences. This provides us with the opportunity to use *ROCK* as a “lab” model of resting and flight height preferences, that can be compared to a wildtype strain. This wildtype (*MID*) *A. aegypti* mosquito strain was derived from mixing wild-caught eggs collected from two *colonias* (i.e. neighborhoods) – colonia Miguel Higalgo (northwest Merida) and colonia Emiliano Zapata Sur (southwest Merida), which together were subsequently reared at UCBE.

In order to avoid cross contamination in the laboratory during the rearing process, *A. aegypti* *ROCK* eggs were physically separated from the *MID* mix of eggs. F1 or F2 larvae from each strain were reared in larval trays at a density of 400-600 larvae/tray and provided with food *ad-libitum* daily to establish the colonies that provided adults for the experiments. The feeding solution provided to the larvae was made by diluting sifted liver powder in reverse osmosis (RO) water to a final concentration of 0.01 gr/ml. After molting to the pupae stage, individuals were placed in emergence cups inside mosquito cages (Bugdorms) in a manner that would result in a density of 400-600 adult mosquitoes/cage. After emerging from the pupae, adult mosquitoes were provided with a 10% sucrose solution *ad libitum* and allowed in the cage unbothered for 48 – 72 hours to allow for complete metamorphosis.

After the 48 – 72 hours, adult mosquitoes were picked out at random from the cages and separated according to sex in order to be used for the hut experiments. In the case of males, each subgroup contained 100 individuals. Females were further distinguished by blood-fed status, with one subgroup containing 100 females that were kept unfed and a second subgroup of 100 females that were blood-fed by arm 2 hours prior. In the case of all groups, the sucrose solution was removed from the cages at 3:00PM local time a day before their respective experiments. All mosquitoes were reared at 27°C, 60% relative humidity and in a 12dark:12light hour photo period.

#### Experimental procedures

The interior of the experimental hut was cleaned a week prior to the start of any experiments, to ensure no spider webs were present, and that the walls were in good condition to set up the sticky traps. The next Monday, four fully-black sticky traps and four HOBO loggers were set up within the hut using heavy-duty double-sided tape. Experiments began the next day (Tuesday) and continued on a daily basis (excluding Sundays) through the next four weeks, as described.

In brief, experiments followed the following protocol. In the morning (~8:30AM) The cover of the cutout on the front wall of the hut was removed and the PFMD placed on top of the metal baseboard in a position to ensure a tight seal. The PFMD’s power supply and monitor were plugged into a surge protector, which was then plugged into an outlet just behind the experimental hut. The monitor’s HDMI chord was plugged into the back of the PFMD unit, as well as the keyboard and mouse. With everything plugged in, the PFMD was then turned on by pushing its power button. To ensure proper recording conditions, the date and time settings of the PFMD were manually set to the nearest second. To make sure the brightness settings of each of the cameras of the PFMD were optimized, they were also manually adjusted as described previously. At this point, the “Run Timed” recording option was selected, set to record for 120 minutes, and given the code name of the upcoming replicate.

With the recording settings all set to go, the group of mosquitoes to be assessed first (i.e. at 9:30AM) were brought into the hut in a container (comprised of a 1.5L plastic container, with a mesh that was strapped to the top of the container using a rubber band) at ~9:10AM and given a 20-minute climatization period. After the 20-minutes, mosquitoes were released from the outside using a string attached to a cotton ball that was placed inside of the mesh covering the container. The string was run through a hook that was attached to the ceiling of the hut (using a command strip) and out the front wall of the hut through a mosquito release sleeve that had been taped shut. Given the PFMD’s blind spot described above, when the cotton ball with the mesh were pulled to the ceiling of the hut, they were outside of the PFMD’s active region and thus did not pose any issues during the recording. Right when the mesh reached the ceiling of the hut, the PFMD’s “Run Timed” setting was clicked to begin recording.

After 120 minutes, one individual swiftly entered the hut with a mesh and a Prokopack aspirator. The mesh was used to cover the release container to prevent mosquitoes who stayed inside of it from flying into the hut. The Prokopack was then used to aspirate all individuals that did not land on any of the sticky traps during the time of the experiment. Aspiration was done for a period of 10-minutes and in a manner that would prevent, as much as possible, any of the flying mosquitoes from getting caught. After the aspiration time, the Prokopack’s collection cup and the covered release container were then taken and placed inside of a -4 C freezer to knockdown the mosquitoes for counting later on. Returning back to the hut, a tape measure was used in order to measure the resting heights of each mosquito that was caught on all four sticky traps. The tape measure was placed just along side each sticky trap, which was inspected for mosquitoes using a flashlight. When a mosquito was identified, its resting height was measured within +/- 1cm to the nearest tape measure mark. This was repeated for each trap and each caught mosquito. Once all caught mosquitoes were measured, the mosquitoes that were in the release container and Prokopack collection cup were counted and summed to ensure all 100 released mosquitoes were accounted for. In the case that the sum did not add to 100, a second 10-minute round of aspiration was done (or until all missing individuals were accounted for, though there were instances where this was not possible). Afterwards, the caught mosquitoes were removed from the traps, and the noon group was brought into the hut and given its corresponding 20-minute climatization period. After the 20-minutes (~12:30PM), the next recording period was started, and the process just described repeated. The same was done for the afternoon group (~4PM), and this entire procedure was followed each day through the rest of the week. The only adjustment that was made after the experiments began involved the switching up of the sticky traps on a weekly basis, as described previously.

#### Statistical analyses

**PFMD Flight Data**

Within the PFMD web-server, a filter was applied to only include mosquito tracks that 1) were longer than one second and 2) spatially occurred beyond 400cm in depth (i.e. z-axis > 400cm). This filter ensured the highest quality track data was used in the analyses given that beyond 400cm is the most complete area of the PFMD’s active region (the square pyramid mentioned previously). With this filter applied, all CSV files corresponding to each of the 72 replicates were exported from the PFMD cloud-server and downloaded to a local PC. Absolute zero was set for each replicate by taking the minimum x-coordinate value and adding it to all x-coordinates in the file; the same process was done for the y-coordinates.

The coordinate sets (x,y,z) of all replicates were first pooled according to strain (i.e. Lab, wt) and visualized by estimating density distributions of each for observed height coordinates (y-coordinates). Three kinds of Generalized Linear Mixed Models (GLMM) were used to analyze the data: 1) fixed effects only included endogenous factors (strain and sex/physiological status), 2) fixed effects only included exogenous factors (color and time of day), and 3) fixed effects included both endogenous and exogenous factors. A “global” model for each kind above was coded and then dredged, to determine which combination of factors and interaction terms provided the best fitting model given the data. Follow up multi-model selection analyses were done by comparing AIC and Delta values. All models used a Beta link function, with “replicate” as the only random effect. The Beta function was picked because all possible values of the y-coordinates were confined within zero to 2-meters; because this distribution is defined on the interval [0,1] all y-coordinates were normalized (i.e. divided by the total height of the hut). In order to link the microclimatic data (mean temperature and mean relative humidity) to each y-coordinate observed, a table join was performed with the HOBO data using “datetime” and respective height. The midpoint between each possible pair of HOBOs was determined. If a y-coordinate had a value above a midpoint, it was assigned the microclimatic data of the HOBO that was above that midpoint; alternatively, if a y-coordinate had a value below a midpoint, it was assigned the microclimatic data of the HOBO that was below the midpoint. Given the high correlation between our microclimatic variables, we decided to not include them directly, but rather as a proxy using time of day.

**Sticky Trap Resting Data**

In a very similar manner to the PFMD track data, we analyzed the sticky trap resting data. Three kinds of GLMMs (endogenous only, exogenous only, and both together) were used to further assess the individual and joint effects of strain, sex/physiological state, color treatment, time of day (proxy for microclimate) on resting height. A “global” model for each kind above was coded and then dredged, to determine which combination of factors and interaction terms provided the best fitting model given the data. Follow up multi-model selection analyses were done by comparing AIC and Delta values. Again, all models used a Beta link function, with “replicate” fit as the only random effect. The Beta function was picked because all possible values of resting height were confined within zero to 2-meters; because this distribution is defined on the interval [0,1] all values of resting height were normalized (i.e. divided by the total height of the hut). In order to link the microclimatic data (mean temperature and mean relative humidity) to each resting height, the measurements of each HOBO (i.e. at each of the four set heights) were pooled and their means calculated. The midpoint between each possible pair of HOBOs was determined. If a mosquito had a resting height above a midpoint, it was assigned the microclimatic data of the HOBO that was above the midpoint; alternatively, if a mosquito had a resting height below a midpoint, it was assigned the microclimatic data of the HOBO that was below the midpoint. Given the high correlation between our microclimatic variables, we decided to not include them directly, but rather as a proxy using time of day.

All analyses presented here were performed using R (version 4.3.2) within R-Studio (version 2023.6.0.421). The R packages used for these analyses were *glmmTMB* (version 1.1.8), *MuMIn* (version 1.47.5), and *ggeffects* (version 1.5.2).

### Supplementary Results

#### Modeling Ae. aegypti innate behavior

Using the model described in Methods section, we simulated N mosquito tracks based on the empirical distribution of track length and turning angles. We initialized simulation with a random set of XYZ coordinates and random heading in three-dimensional space. To simulate a subsequent step, we then pulled from each theoretical turning angle distribution to update the heading along the XY and XZ planes and pulled from the theoretical step-length distribution to draw a movement distance, with the new XYZ coordinates calculated from these metrics in relation to the initial point of our simulated individual. We note that this assumes that the step length and both turning angle distributions are independent. Repeating this process a number of times allowed the generation of a null random walk trajectory emulating mosquito movement in the absence of persistency or bias in their movement. Emulated trajectories were produced for each experimental group (unfed females, blood-fed females, and males), and together with all observed trajectories, assessed using Generalized Linear Mixed Models (GLMMs) to detangle the fixed effects of endogenous factors (strain and sex/physiological status). A “global” model including all factors (strain, sex, and their interaction) was initially fitted, and subsequently dredged (*MuMin* R package) to obtain a table of all possible models for subsequent model selection using AICc (Table S1). A similar procedure was followed for GLMMs comparing between the Lab-strain trajectories and the wt-trajectories. Given both of these trajectory results, the GLMM corresponding to resting height was only fitted to the Unfed female trajectories from both the Lab and wt strains.

**Table S1.** Comparative statistics for all possible models when only considering endogenous factors (strain, sex/physiological status, and their interaction). Models were fit independently for flight behavior (PFMD Y-coordinates) and resting behavior (sticky trap measurements). All models contained replicate as a random effect (1|replicate.id).

| **FLIGHT (comparing Simulated, Lab-strain, and wt-strain trajectories)** | | | | | | | | | | |
| --- | --- | --- | --- | --- | --- | --- | --- | --- | --- | --- |
| **MODEL** | **Conditional**  **Intercept** | **Dispersion**  **Intercept** | **Sex/**  **Phys** | **strain** | **sex:strain** | **df** | **LogLik** | **AICc** | **delta** | **weight** |
| ~ strain * sex | 1.818 | **+** | **+** | **+** | **+** | 11 | 1,216,883 | -2,433,743 | 0 | 0.59 |
| ~ strain + sex | 1.900 | **+** | **+** | **+** | - | 7 | 1,216,878 | -2,433,742 | 0.733 | 0.41 |
| ~ strain | 1.734 | **+** | - | **+** | - | 5 | 1,216,867 | -2,433,723 | 19.76 | 3.02 x 10^-5^ |
| Null | -0.919 | **+** | - | - | - | 3 | 1,216,841 | -2,433,676 | 67.46 | 1.32 x 10^-15^ |
| ~ sex | -0.773 | **+** | **+** | - | - | 5 | 12,070,898 | -24,141,787 | 8.27 | 0.01 |
| **FLIGHT (comparing Lab-strain and wt-strain trajectories)** | | | | | | | | | | |
| **MODEL** | **Conditional**  **Intercept** | **Dispersion**  **Intercept** | **Sex/**  **Phys** | **strain** | **sex:strain** | **df** | **LogLik** | **AICc** | **delta** | **weight** |
| ~ strain * sex | -0.564 | **+** | **+** | **+** | **+** | 8 | 3,045,657 | -6,091,298 | 0 | 0.69 |
| ~ strain + sex | -0.578 | **+** | **+** | **+** | - | 6 | 3,045,654 | -6,091,296 | 1.753 | 0.29 |
| ~ sex | -0.736 | **+** | **+** | - | - | 5 | 3,045,651 | -6,091,291 | 6.526 | 0.026 |
| Null | -0.940 | **+** | - | - | - | 3 | 3,045,643 | -6,091,281 | 17.20 | 1.2 x 10^-4^ |
| ~ strain | -0.820 | **+** | - | **+** | - | 4 | 3,045,644 | -6,091,280 | 17.60 | 1.04 x 10^-4^ |
| **RESTING (comparing Lab-strain and wt-strain)** | | | | | | | | | | |
| **MODEL** | **Conditional**  **Intercept** | **Dispersion**  **Intercept** | **Sex/ Phys** | **strain** | **sex:strain** | **df** | **LogLik** | **AICc** | **delta** | **weight** |
| ~ strain | -0.208 | **+** | - | **+** | - | 4 | 29.761 | -51.216 | 0 | 0.507 |
| ~ strain + sex | -0.185 | **+** | **+** | **+** | - | 6 | 31.797 | -50.943 | 0.272 | 0.443 |
| ~ strain * sex | -0.1872 | **+** | **+** | **+** | **+** | 8 | 31.817 | -46.501 | 4.715 | 0.048 |
| Null | -0.531 | **+** | - | - | - | 3 | 22.629 | -39.076 | 12.139 | 0.001 |
| ~ sex | -0.599 | **+** | **+** | - | - | 5 | 24.543 | -38.625 | 12.591 | 0.0001 |

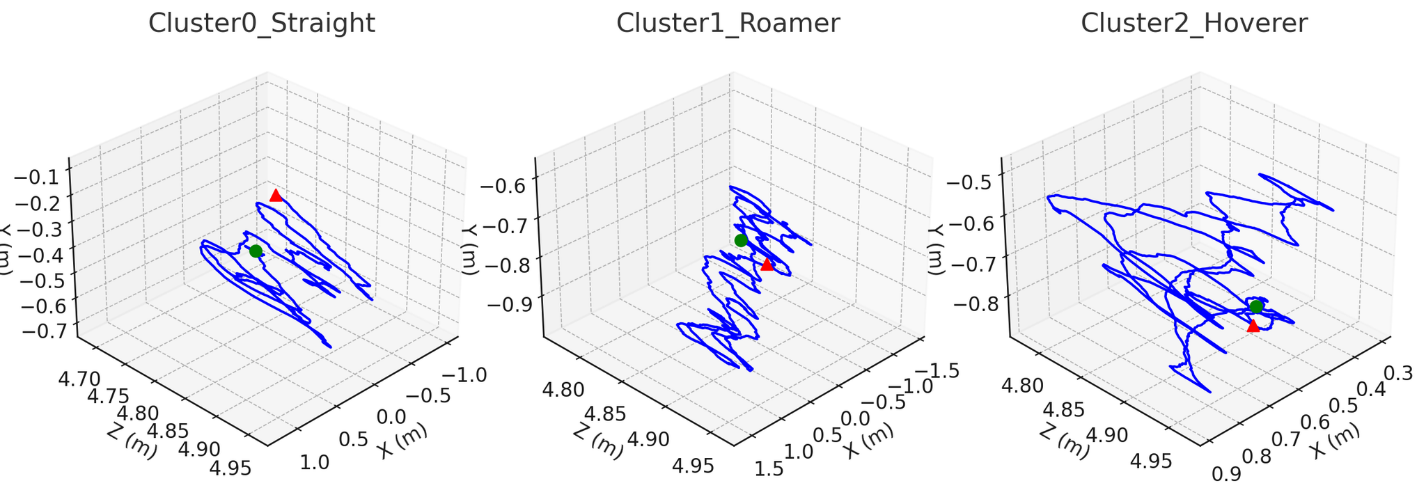

**Figure S4.** 3D scatterplots comparing two different mosquito tracks based on their estimated fractal dimension (D) to highlight the difference between a trajectory with fewer turns and thus more linear (left plot) vs a trajectory with many turns and thus tortuous/exploratory (right plot).

#### Temperature and relative humidity

Pooling all microclimate data according to HOBO (see ‘Experimental Set Up’ above) revealed marked gradients for both temperature and relative humidity from the floor to the ceiling of the hut (Fig. S5). Mean temperatures were significantly different across all possible logger pairs and demonstrated a strong positive relationship with height, with the strongest difference occuring between the hottest temperatures at the ceiling versus the coolest temperatures at the floor (Fig. S5). A similar, but opposite pattern was observed for RH with the highest humidities at the floor versus the lowest humidities at the ceiling (Fig. S5). A Pearson’s Correlation test showed a significant negative correlation of -0.914 between all temperature and relative humidity measurements (Fig. S5).

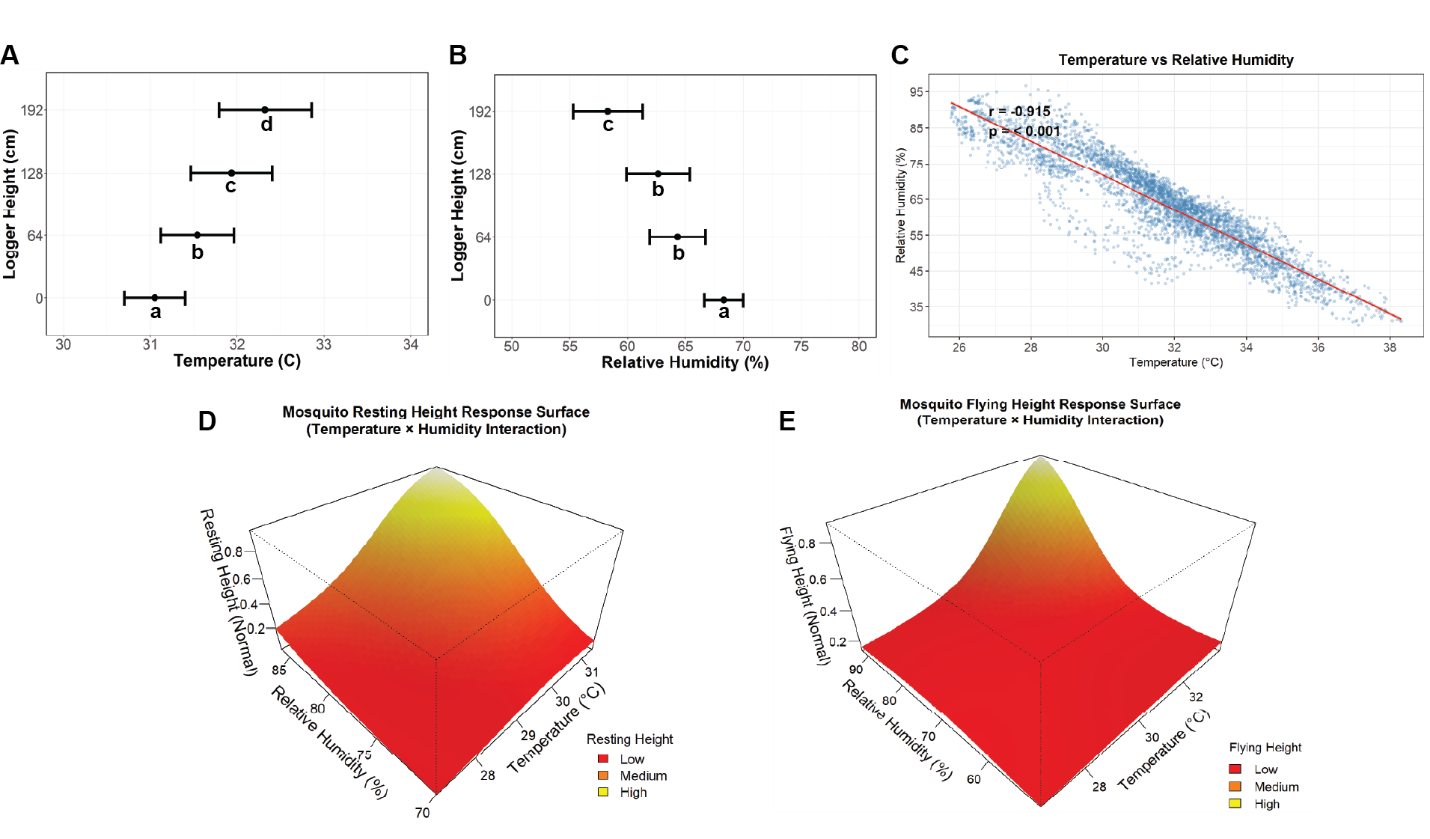

**Figure S5. Summary of microclimatic parameters and their significant correlation. A** Plot depicting the mean temperature in Celsius (points) and corresponding CI_95%_ (error bars) for each logger at their respective height (i.e. 0cm, 64cm, 128cm, and 192cm). Different letters indicate significance between loggers. **B** Plot depicting the mean relative humidity in percent (points) and corresponding CI_95%_ (error bars) for each logger at their respective height (i.e. 0cm, 64cm, 128cm, and 192cm). Different letters indicate statistical significance between loggers, with the same letters indicating no significance. **C** Scatterplot showing the strong negative correlation between temperature and relative humidity across all microclimatic measurements. The red represents a linear regression fit (r = -0.915, *p* < 0.001).

The 3D visualizations of the GAM we fitted to detangle the relationship between temperature, relative humidity, and height are quite similar for both resting and flying height of wt mosquitoes. Under high temperatures (> 29 C) and high relative humidities (> 75%), the tensor surfaces estimate a maximum peak of 2 meters for both flight and resting (brightest yellow); Figure 2A-B). When temperatures are low (< 29), the tensor surfaces show mosquitoes resting and flying at the lowest heights (bright red) regardless of the relative humidity percentage; Figure 2A-B). However, if temperatures are high (> 29 C) mosquitoes appear to only fly and rest at low heights if relative humidities are below 75% Thus it appears that at high temperatures mosquitoes prefer to fly and rest at low heights, but an increase in relative humidity can offset this preference and instead lead mosquitoes to fly and rest at higher heights. In recent times, multiple authors have highlighted the idea that mosquitoes are likely to move through space following moisture gradients^3^. At the microclimatic level, we show that air temperature and humidity, but more importantly humidity, may be driving the innate behavior of Ae. aegypti.

To further investigate the collinearity of temperature and relative humidity and their combined effect on resting height, we fitted additional Generalized Additive Models (GAMs). Two kinds of models were built – 1) mean temperature and mean relative humidity as smoothing terms and 2) mean temperature and mean relative humidity as an interaction term (Table S2). Both of these models were fitted with the Beta distribution as the link function, and set to use the Restricted Maximum Likelihood (REML) method for parameter estimation. Regardless of which type of model, it remained quite difficult to separate the effect of temperature and humidity on mosquito behavior. Thus, we decided to calculate vapor pressure deficit (VPD; Fig. S6), which is considered to be more relevant for mosquito physiological and ecological traits^4–6^. VPD is defined as the difference between the saturation vapor pressure at a given temperature (es) and the actual vapor pressure (ea):

VPD = es - ea = (0.6108 × exp((17.27T) / (T + 237.3))) - ((RH / 100) × es)

where T is air temperature (°C) and RH is relative humidity (%). Particularly for mosquitoes, lower VPD values indicate more favorable microclimates for flight and resting, as desiccation risk is reduced. GAMs using VPD were contrasted to the ones using temperature and relative humidity (Table S2). The R packages used for these analyses were *mgcv* (version 1.9.0) and *rgl* (version 1.2.8).

Lastly, we modeled the probability of *Ae. aegypti* to fly and rest above 1 meter as a function of VPD and time of day using a binomial generalized linear model (GLM) with a logit link. To address issues of small-sample bias in maximum likelihood estimation, the model was fit using the Firth method implemented via median bias-reducing adjusted score (AS-median) equations in the *brglm2* package in R.

**Table S2.** Comparative statistics of all Generalized Additive Models (GAMs) used to further investigate the relationship between temperature, relative humidity, VPD, and observed mosquito flight/resting height. Models were fit independently for flight behavior (PFMD Y-coordinates) and resting behavior (sticky trap measurements). Models were fitted to data from wt unfed females assessed under no attraction (i.e. White sticky traps).

| **FLIGHT** | | | | | | | |
| --- | --- | --- | --- | --- | --- | --- | --- |
| **MODEL** | **Terms** | **edf** | **Chi-sq** | **p-value** | **AIC** | **Deviance Explained** | **REML** |
| y-coordinate ~  te(mean_temp, mean_rh,  k = 3) | te(Temp., RH) | 8 | 49,029 | **< 0.0001** | -165,339 | 45.8% | -82,633 |
| y-coordinate ~  s(mean_temp, k = 3) | s(Temp) | 2 | 19,007 | **< 0.0001** | -142,628 | 22.3% | -71,302 |
| y-coordinate ~ s(mean_rh, k = 3) | s(RH) | 2 | 19,267 | **< 0.0001** | -143,316 | 23.2% | -71,646 |
| y-coordinate ~  s(VPD, k = 3) | s(VPD) | 2 | 21,194 | **< 0.0001** | -145,067 | 25.3% | -72,521 |
| **RESTING** | | | | | | | |
| **MODEL** | **Terms** | **edf** | **Chi-sq** | **p-value** | **AIC** | **Deviance Explained** | **REML** |
| Resting height ~ te(mean_temp, mean_rh,  k = 3) | te(Temp., RH) | 3 | 34.56 | **< 0.0001** | -35.40 | 79.4% | -25.59 |
| Restinh height ~  s(mean_temp, k = 3) | s(Temp) | 1.69 | 2.195 | 0.241 | -19.96 | 25.6% | -11.53 |
| Resting height ~ s(mean_rh, k = 3) | s(RH) | 1 | 2.391 | 0.122 | -20.83 | 19.9% | -11.97 |
| Resting height ~ s(VPD, k = 3) | s(VPD) | 1 | 2.128 | 0.145 | -20.55 | 18.3% | -11.84 |

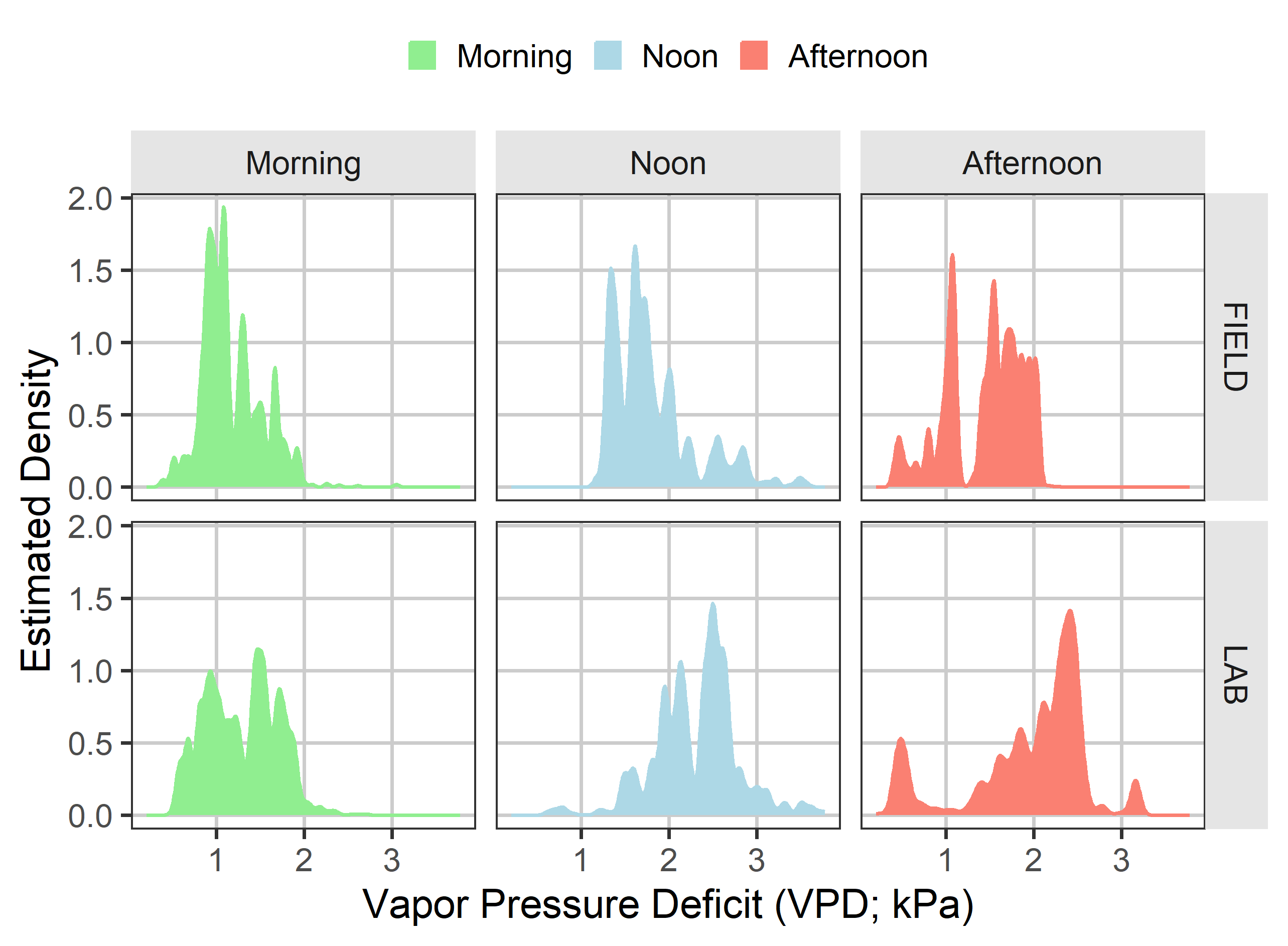

**Figure S6. Density curves portraying the distribution of pooled VPD estimates stratified by Strain (wt – top row; Lab – bottom row) and Time of Day (morning in green, noon in blue, and afternoon in red).** Mosquitoes from both strains appear to have been exposed to the same range of VPD values (0 to 3 kPa), with Morning and Noon distributions having most overlap, while Afternoon distributions showed the most heterogeneity. This would make sense given the experiments were done during the wet season, with heavy rain like conditions occurring mainly in the Afternoon hours (2pm – 5pm local time).

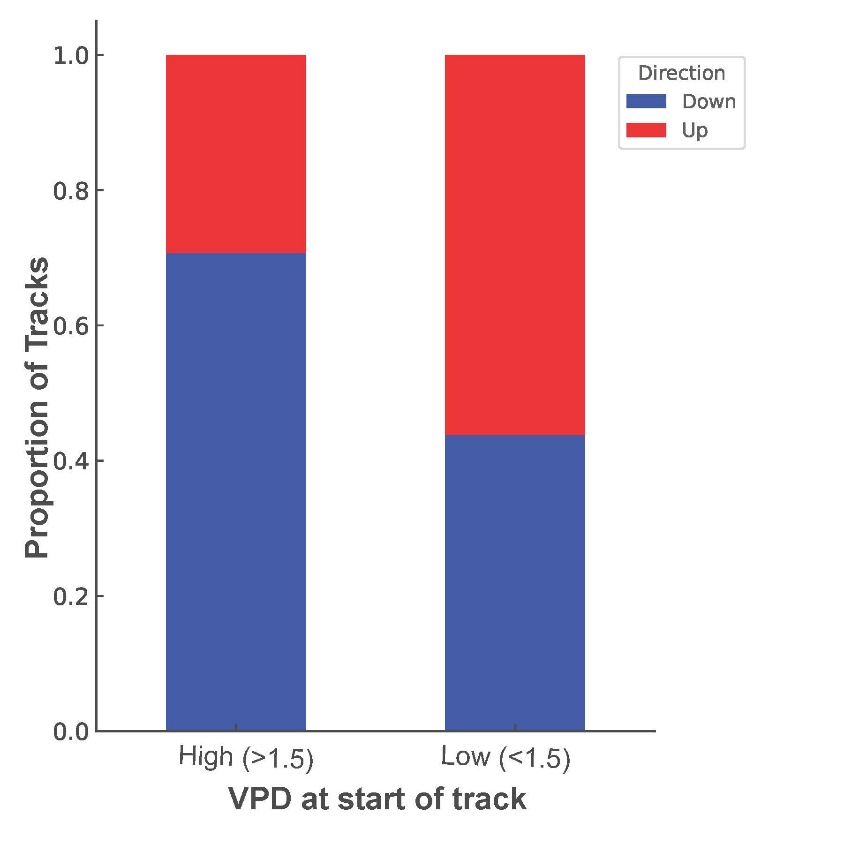

**Figure S7. Relationship between VPD at the beginning of the track and the tendency of the mosquito track to end at a higher (up) or lower (down) point.** Result using only tracks with displacement higher than 50cm (total of 318 tracks,), and using wt mosquitoes and white color background (no stimuli). When mosquito tracks started at VPD above 1.5, 71% of the tracks moved towards lower heights. However, when VPD at the beginning of the track was below 1.5 kPA only 44 mosquitoes moved low. The Odds of flying low at high VPD, tested with a binomial GLM, was 3.1 (95%CI = 1.95-4.94).

#### Resting Color

Given we were unable to detangle the relationship between our microclimatic metrics (temperature and relative humidity), in addition to the counterintuitive result we obtained when using VPD as an index for desiccation pressure, we decided to use Time of Day as a proxy for different microclimatic profiles (Fig. S6). As such, Time of Day was included as a fixed effect in all exogenous GLMMs.

**Table S3.** Comparative statistic results for all possible models (dredged from global model) when only considering exogenous factors (color and time of day). Models were fit independently for flight behavior (PFMD Y-coordinates) and resting behavior (sticky trap measurements). All models contained replicate as a random effect (1|replicate.id).

| **FLIGHT** | | | | | | | | | |
| --- | --- | --- | --- | --- | --- | --- | --- | --- | --- |
| **MODEL** | **Conditional**  **Intercept** | **Dispersion**  **Intercept** | **color** | **time of day** | **df** | **LogLik** | **AICc** | **delta** | **weight** |
| ~ color | -1.374 | **+** | **+** | - | 6 | 2,619,186 | -5,238,360 | 0 | 0.714 |
| ~ color + time of day | -1.337 | **+** | **+** | **+** | 8 | 2,619,187 | -5,238,358 | 1.83 | 0.285 |
| Null | -1.180 | **+** | - | - | 3 | 2,619,173 | -5,238,340 | 19.73 | 3.7 x 10^5^ |
| ~ time of day | -1.144 | **+** | - | **+** | 5 | 2,619,173 | -5,238,336 | 23.50 | 5.6 x 10^-6^ |
| **RESTING** | | | | | | | | | |
| **MODEL** | **Conditional**  **Intercept** | **Dispersion**  **Intercept** | **color** | **time of day** | **df** | **LogLik** | **AICc** | **delta** | **weight** |
| ~ color | -1.158 | **+** | **+** | - | 6 | 61.16 | -109.91 | 0 | 0.87 |
| ~ color + time of day | -1.253 | **+** | **+** | **+** | 8 | 61.40 | -106.09 | 3.82 | 0.13 |
| Null | -0.598 | **+** | - | - | 3 | 46.66 | -87.21 | 22.7 | 1 x 10^-5^ |
| ~ time of day | -0.51 | **+** | - | **+** | 5 | 45.74 | -83.203 | 26.7 | 1.4 x 10^-6^ |

#### Flight and resting patterns between sexes and physiological states

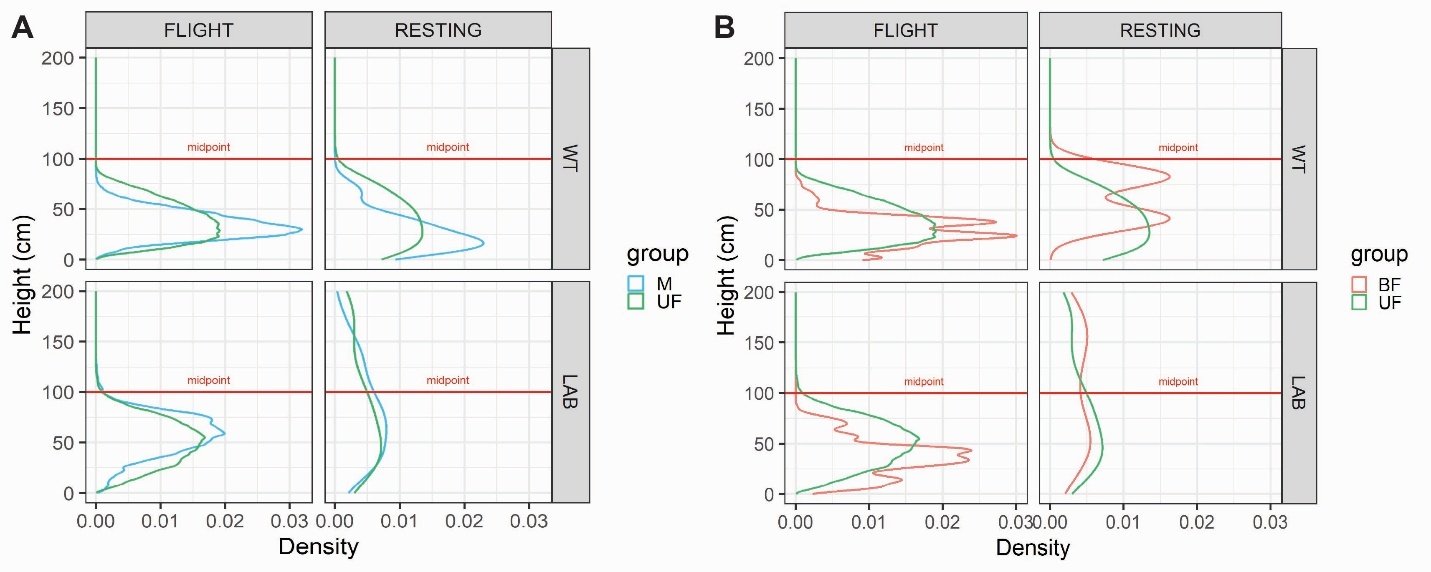

**Figure S8**. Estimated density distributions comparing the flight and resting behavior of Males vs Unfed Females (A – left) and of Unfed Females vs Blood-Fed Females (B – right). Each panel (i.e. both A and B) was further stratified according to behavior (i.e. Flight on the left and Resting on the right) and strain (wt on the top row and Lab on the bottom row). The solid red line represents the vertical midpoint inside the hut. Note that the Y-axis corresponds to Height (cm) and the X-axis to the estimated density of mosquitoes recorded/caught at all possible heights between 0 and 200cm.

**Table S4. Sex and physiological status as drivers for *Aedes aegypti* resting height.** Summarizing results of linear mixed effects model fitting the fixed effect of strain, sex/physiological status, and their interaction on observed resting height. Models were fit to the data corresponding to replicates without color attraction (i.e. White sticky traps) and had replicate coded as the only random effect (1|replicate.id). Note that “Lab-adapted” was set as the reference level for strain type, and “Unfed Females” as the reference sex and/or physiological state.

| **Trait** | **Factor** | **Coeff (95%CI)** | **z** | **p-value** |
| --- | --- | --- | --- | --- |
| **Resting^1^**  (~ strain [Lab : wt] * sex) | Intercept | -0.187 (-0.573, 0.198) | -0.951 | 0.341 |
|  | **Strain – wt** | **-1.01 (-1.679, -0.332)** | **-2.925** | **0.0035 **** |
|  | Sex – Blood-Fed | 0.471 (-0.279, 1.220) | 1.232 | 0.218 |
|  | Sex – Male | 0.163 (-0.655, 0.328) | -0.652 | 0.515 |
|  | Strain – wt : Sex – Blood-Fed | 0.131 (-1.50, 1.760) | 0.158 | 0.875 |
|  | Strain – wt : Sex – Males | -0.032 (-0.924, 0.859) | -0.071 | 0.943 |
| ^1^ Random Effect Parameters: variance = 0.0293 and standard deviation = 0.1711 (0.053, 0.557) | | | | |

**Table S5*.* Endogenous and exogenous together better explain *Aedes aegypti* behavior.** Summarizing results of linear mixed effects model fitting including the fixed effect of endogenous factors (strain, sex/physiological status, and their interaction) together with exogenous factors (color treatment and time of day). Like the other models, replicate was coded as the only random effect (1|replicate.id). Note that “Lab-adapted” was set as the reference level for strain type, “Unfed Females” as the reference sex and/or physiological state, “White” as the reference color treatment, and “Morning” as the reference time of day.

| **Trait** | **Factor** | **Coeff (95%CI)** | **z** | **p-value** |
| --- | --- | --- | --- | --- |
| **Flight^1^**  (~ strain [Lab : wt] * sex + color + time of day) | Intercept | **-1.056 (-1.341, -0.771)** | **-7.260** | **3.88 x 10^-13^ ***** |
|  | **Strain – wt** | **-0.360 (-0.651, -0.068)** | **-2.420** | **0.0155 *** |
|  | Sex – Blood-Fed | -0.0163 (-0.310, 0.278) | -0.108 | 0.9137 |
|  | Sex – Male | -0.103 (-0.395, 0.189) | -0.692 | 0.4886 |
|  | Strain – wt : Sex – Blood-Fed | 0.0933 (-0.326, 0.513) | 0.436 | 0.6629 |
|  | Strain – wt : Sex – Males | -0.0668 (-0.479, 0.345) | -0.318 | 0.751 |
|  | **Color – Black** | **0.820 (0.577, 1.064)** | **6.609** | **3.87 x 10^-11^ ***** |
|  | **Color – Black:White** | **0.496 (0.253, 0.740)** | **3.998** | **6.39 x 10^-5^ ***** |
|  | Color – White:Black | 0.107 (-0.136, 0.351) | 0.863 | 0.388 |
|  | Time of day – Afternoon | -0.096 (-0.305, 0.114) | -0.896 | 0.370 |
|  | **Time of day - Noon** | **-0.212 (-0.429, -0.01)** | **-2.057** | **0.0397 *** |
| **Resting^2^**  (~ strain [Lab : wt] * sex + color + time of day) | Intercept | -0.239 (-0.533, 0.055) | -1.592 | 0.1114 |
|  | **Strain – wt** | **-1.01 (-0.845, -0.308)** | **-4.207** | **2.58 x 10^-5^ ***** |
|  | Sex – Blood-Fed | 0.471 (-0.174, 0.363) | 0.688 | 0.491 |
|  | Sex – Male | 0.163 (-0.366, 0.093) | -1.166 | 0.244 |
|  | Strain – wt : Sex – Blood-Fed | 0.131 (-0.244, 0.560) | 0.769 | 0.442 |
|  | Strain – wt : Sex – Males | -0.032 (-0.385, 0.342) | -0.115 | 0.908 |
|  | Color – Black | 0.219 (-0.043, 0.482) | 1.639 | 0.1012 |
|  | **Color – Black:White** | **1.055 (0.785, 1.325)** | **7.645** | **2.10 x 10^-14^ ***** |
|  | **Color – White:Black** | **-0.304 (-0.57, -0.038)** | **-2.240** | **0.0251 *** |
|  | Time of day – Afternoon | 0.005 (-0.181, 0.192) | 0.055 | 0.956 |
|  | Time of day - Noon | -0.043 (-0.230, 0.144) | -0.451 | 0.652 |
| ^1^ Random Effect Parameters: variance = 0.1327 and standard deviation = 0.3643 (0.307, 0.431)  ^2^ Random Effect Parameters: variance = 0.0575 and standard deviation = 0.24 (0.184, 0.313) | | | | |

#### *Aedes aegypti* reciprocal crosses and observed resting preferences

**Table S6*.* Reciprocal cross progeny resemble resting preferences of wt parents.** Summarizing results of linear mixed effects models comparing observed resting heights between cross progeny and their respective parents. Like the other models, replicate was coded as the only random effect (1|replicate.id). Note that “Progeny” was set as the reference level for strain type.

| **Cross** | **Factor** | **Coeff (95%CI)** | **z** | **p-value** |
| --- | --- | --- | --- | --- |
| **Lab Male x wt Female^1^**  ~ strain [each parent : Progeny | **Intercept** | **-0.606 (-0.840, -0.373)** | **-5.094** | **3.51 x 10^-7^ ***** |
|  | **Strain – Lab Male** | **0.394 (0.039, 0.748)** | **2.176** | **0.0296 *** |
|  | Strain – wt Female | -0.0001 (-0.3863, 0.3860) | -0.001 | 0.9994 |
| **Lab Female x wt Male^2^**  ~ strain [each parent : Progeny] | **Intercept** | **-0.494 (-0.686, -0.302)** | **-5.038** | **4.7 x 10^-7^ ***** |
|  | **Strain – Lab Female** | **0.516 (0.231, 0.801)** | **3.546** | **3.91 x 10^-4^ ***** |
|  | Strain – wt Male | -0.121 (0.053, 0.356) | -0.737 | 0.461 |
| ^1^ Random Effect Parameters: variance = 0.044 and standard deviation = 0.21 (0.118, 0.373)  ^2^ Random Effect Parameters: variance = 0.0188 and standard deviation = 0.137 (0.053, 0.356) | | | | |
